# A comparison of one-rate and two-rate inference frameworks for site-specific *dN/dS* estimation

**DOI:** 10.1101/032805

**Authors:** Stephanie J. Spielman, Suyang Wan, Claus O. Wilke

**Author notes:** Current Address: Institute for Genomics and Evolutionary Medicine, Temple University, Philadelphia, PA 19122, USA.

## Abstract

Two broad paradigms exist for inferring *dN/dS*, the ratio of nonsynonymous to synonymous substitution rates, from coding sequences: i) a one-rate approach, where *dN/dS* is represented with a single parameter, or ii) a two-rate approach, where dN and dS are estimated separately. The performances of these two approaches have been well-studied in the specific context of proper model specification, i.e. when the inference model matches the simulation model. By contrast, the relative performances of one-rate vs. two-rate parameterizations when applied to data generated according to a different mechanism remains unclear. Here, we compare the relative merits of one-rate and two-rate approaches in the specific context of model misspecification by simulating alignments with mutation-selection models rather than with *dN/dS*-based models. We find that one-rate frameworks generally infer more accurate *dN/dS* point estimates, even when dS varies among sites. In other words, modeling *dS* variation may substantially reduce accuracy of *dN/dS* point estimates. These results appear to depend on the selective constraint operating at a given site. In particular, for sites under strong purifying selection (*dN/dS*<~0.3), one-rate and two-rate models show comparable performances. However, one-rate models significantly outperform two-rate models for sites under moderate-to-weak purifying selection. We attribute this distinction to the fact that, for these more quickly evolving sites, a given substitution is more likely to be nonsynonymous than synonymous. The data will therefore be relatively enriched for nonsynonymous changes, and modeling dS contributes excessive noise to *dN/dS* estimates. We additionally find that high levels of divergence among sequences, rather than the number of sequences in the alignment, are more critical for obtaining precise point estimates.

## Introduction

A variety of computational approaches have been developed to infer selection pressure from protein-coding sequences in a phylogenetically-aware context. Among the most commonly used methods are those which compute the evolutionary rate ratio *dN/dS*, which represents the ratio of non-synonymous to synonymous substitution rates. Beginning in the mid-1990s, this value has been calculated using maximum-likelihood (ML) approaches (Goldman and Yang 1994; Muse and Gaut 1994), and since then, a wide variety of inference frameworks have been developed to infer *dN/dS* at individual sites in protein-coding sequences (Nielsen and Yang 1998; Yang *et al.* 2000; Yang and Nielsen 2002; Yang and Swanson 2002; Kosakovsky Pond and Frost 2005; Kosakovsky Pond and Muse 2005; Murrell *et al.* 2012c; Lemey *et al.* 2012; Murrell *et al.* 2013).

Typically, the performance of evolutionary models is examined using simulation-based approaches wherein sequences are simulated according to the model being examined, and inferences are subsequently performed on the simulated data. This approach ensures that simulated and inferred parameters correspond. Although useful, this strategy fundamentally assumes that the data being analyzed was generated by the same mechanism that the inference model used, and hence the model has been correctly specified. Crucially, this scenario does not apply to the analysis of natural sequence data. Indeed, real genomes evolve according to a variety of interacting evolutionary forces, not according to a phylogenetic model of sequence evolution. As such, it remains unclear how well evolutionary models perform when they are applied to sequence data that has been technically *misspecified*, i.e. where the data does not conform to the inference model’s assumptions. Indeed, ML-based inferences methods are guaranteed to converge upon the true parameter value when the model is properly specified (provided there is sufficient data), but there is no such guarantee when the data violates critical model assumptions (Yang 2006).

We therefore seek to we extend our understanding of *dN/dS*-based inference model performance by studying how well these models infer site-specific *dN/dS* point estimates when the model is mathematically misspecified. To this end, we simulate sequences not under a *dN/dS*-based model, but instead under the mutation–selection (MutSel) modeling framework. Unlike *dN/dS*-based models, MutSel models are based on population genetics principles and describe the site-specific evolutionary process as a dynamic interplay between mutational and selective forces (Halpern and Bruno 1998; Yang and Nielsen 2008). Therefore, many regard MutSel models as more mechanistically representative of real coding-sequence evolution than *dN/dS*-based models, which are primarily phenomenological in nature (Thorne *et al.* 2007; Holder *et al.* 2008; Rodrigue *et al.* 2010; Thorne *et al.* 2012; Tamuri *et al.* 2012; Liberles *et al.* 2013). Indeed, substitution rate itself is not an evolutionary mechanism, but rather an emergent property of various interacting evolutionary processes. For this reason, MutSel-based simulation approaches have been used to study the behavior of phylogenetic and evolutionary rate inferences (Holder *et al.* 2008; McCandlish *et al.* 2013; Spielman and Wilke 2015b; dos Reis 2015).

Recently, we introduced a mathematical framework which allows us to accurately calculate a *dN/dS* ratio directly from the parameters of a MutSel model (Spielman and Wilke 2015b). This framework gives rise to a robust benchmarking strategy through which we can simulate sequences using a MutSel model, infer *dN/dS* on the simulated sequences using established approaches, and then compare inferred to expected *dN/dS* given the parameters of the MutSel model. We have previously successfully used such an approach to identify biases in *dN/dS* inference approaches for whole-gene evolutionary rates (Spielman and Wilke 2015b). Here, we employ this approach to evaluate the performance of site-specific *dN/dS*-based inference approaches.

Importantly, because the *dN/dS* models used for inference are mathematically misspecified to the data, we can expect inferences to feature statistical biases, which in turn illuminate how these models might behave in more complex scenarios. Several important distinctions between *dN/dS* and MutSel models contribute to the model misspecification examined here. Most importantly, while *dN/dS* models assume that all nonsynonymous substitutions occur at the same rate, MutSel models assume different rates for each type of nonsynonymous substitution, and similarly for synonymous substitutions in more complex scenarios. This distinction gives rise to a key difference in model assumptions. In particular, because the *dN/dS* parameter is a rate constant, *dN/dS* models implicitly assume that substitutions are Poisson-distributed in time. By contrast, in the MutSel model, *dN/dS* is not a rate constant, and thus substitutions will be over-dispersed relative to a Poisson process.

Two primary questions motivate the present study: i) How accurate are various inference frameworks for *dN/dS* point estimation? and ii) Under what conditions does *dN/dS* capture the long-term evolutionary dynamics of site-specific coding-sequence evolution? For the first question, we focus our efforts on distinguishing performance between two *dN/dS* inference paradigms: one-rate and two-rate models. One-rate models parameterize *dN/dS* with a single parameter for *dN*, effectively fixing *dS* = 1 at all sites, whereas two-rate models use separate parameters for *dN* and *dS* at each site. Some studies have suggested that the two-rate paradigm leads to more robust positive-selection inference (Kosakovsky Pond and Muse 2005; Murrell *et al.* 2013), whereas others have suggested that the extra *dS* parameter may actually confound positive selection inference (Yang *et al.* 2005; Wolf *et al.* 2009). Here, we do not benchmark positive-selection inference, but we instead ask how this parameterization affects *dN/dS* point estimation in the context of model misspecification.

The second question arises naturally from our use of MutSel models, which describe the equilibrium site-specific codon fitness values. As a consequence, any *dN/dS* calculated from MutSel model parameters describes, by definition, the steady-state *dN/dS* (Spielman and Wilke 2015b). Since *dN/dS* is an inherently time-sensitive measurement (Rocha *et al.* 2006; Kryazhimskiy and Plotkin 2008; Wolf *et al.* 2009; Mugal *et al.* 2014; Meyer *et al.* 2015), it is not necessarily true that *dN/dS* measured from a given dataset will reflect the equilibrium value. Therefore, our approach additionally enables us to identify the conditions under which site-specific *dN/dS* ratios are expected to reflect the long-term, rather than transient, evolutionary dynamics.

## Materials and Methods

### Derivation of MutSel simulation parameters

We simulated heterogeneous alignments, such that each site evolved according to a distinct distribution of codon state frequencies, according to the HB98 MutSel model (Halpern and Bruno 1998) using Pyvolve (Spielman and Wilke 2015a). The rate matrix for this model is given by

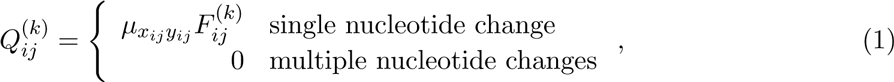

where *µ_x_ij_y_ij__* is the site-invariant mutation rate where *x_ij_* is the focal nucleotide before mutation and *y_ij_* the focal nucleotide after mutation during the substitution from codon *i* to *j*. 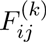 is the fixation probability from codon *i* to *j* at site *k* and is defined as

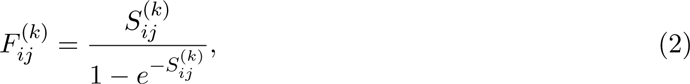

where 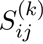 is the scaled selection coefficient from codon *i* to *j* at site *k* (Halpern and Bruno 1998). Thus, this model is specified using a nucleotide-level mutation model (*µ_xy_* parameters) and codon-level fitness values (*F_ij_* parameters).

We simulated four sets of alignments, all of which assumed an HKY85 mutation model (Hasegawa *et al.* 1985) with the transition-transversion bias parameter set to 4. Two alignment sets assumed equal nucleotide frequencies (*π_i_* = 0.25 for *i* ∈ {A, C, G, T}), and the other two alignment sets assumed unequal nucleotide frequencies (arbitrarily set to *π_A_* = 0.32, *π_T_* = 0.28, *π_C_* = 0.18, *π_G_* = 0.22), to incorporate underlying nucleotide compositional bias. We refer to these parameterizations, respectively, as ∏_equal_ and ∏_unequal_. For each mutational parameterization, we simulated an alignment set where all synonymous codons shared the same fitness value (no codon bias) and an alignment set where synonymous codons differed in fitnesses (codon bias). The ∏_equal_ and ∏_unequal_ simulations without codon bias used the same sets of fitness parameters, and similarly the ∏_equal_ and ∏_unequal_ simulations with codon bias used the same sets of fitness parameters.

It has previously been found that site-specific amino acid frequencies, in empirical alignments, tend to follow a Boltzmann distribution (Porto *et al.* 2004; Ramsey *et al.* 2011). Therefore, to derive realistic sets of codon fitnesses for each these simulations, we began by simulating 100 site-specific amino-acid frequency distributions from a Boltzmann distribution:

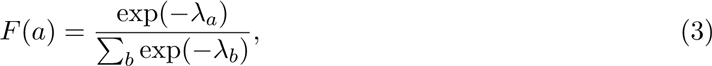

where *F* (*a*) is the state frequency of amino-acid *a*, *a* and *b* index amino acids from 0–19, and the parameter λ increases with evolutionary rate (Ramsey *et al.* 2011). For each frequency distribution, we sampled a value for λ from a uniform distribution 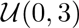, and we selected a random ranking for all amino acids. These frequency calculations formed the basis of our derivation of fitness values used in all simulations.

Importantly, when a symmetric nucleotide mutation model is assumed (e.g. *µ_xy_* = *µ_yx_*), codon fitness values can be calculated directly as the logarithm of codon equilibrium frequency values (Sella and Hirsh 2005). Therefore, we directly computed codon fitness values from the derived frequency values, under the assumption that synonymous codons shared the same fitness. These fitness parameters were employed for both ∏_equal_ and ∏_unequal_ simulations without codon bias.

To derive fitness parameters for simulations with codon bias, we randomly selected a preferred codon for each amino acid. We assigned a state frequency of *γF*(*a*), where γ was drawn from a uniform distribution 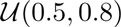, to the preferred codon, and we assigned the remaining frequency (1 − γ)*F*(*a*) evenly to all remaining synonymous codons. In this way, the overall amino-acid state frequency was unchanged, but its synonymous codons occurred with differing frequencies. Note that a single parameter was selected for each frequency distribution (i.e., each resulting alignment position), and not for each set of synonymous codons. Again, fitness distributions were directly computed from these resulting codon frequencies for use in both ∏_equal_ and ∏_uequal_ simulations with codon bias.

Unlike the ∏_equal_ simulations, the ∏_unequal_ simulations did not contain symmetric mutation rates. Therefore, we obtained stationary codon frequencies for the ∏_unequal_ simulations, for use in *dN/dS* calculations, numerically as the dominant eigenvector of each MutSel model’s matrix, which was constructed from mutation rates and codon fitness values. In this way, all stationary codon frequency distributions incorporated, by definition, information regarding both codon-level fitness and nucleotide-level mutation.

We simulated heterogeneous alignments across an array of balanced phylogenies, containing either 128, 256, 512, 1024, or 2048 sequences. For each number of taxa, we simulated sequences with varying degrees of divergence, with all branch lengths equal to either 0.0025, 0.01, 0.04, 0.16, or 0.64. Throughout, we use *N* to refer to a given simulation’s number of taxa and *B* to refer a given simulation’s branch length. We simulated 50 alignment replicates for each combination of *N* and *B*. We additionally simulated alignments, using only the ∏_unequal_ parameterizations, along five different empirical phylogenies (Table 1), again with 50 replicates each. For these simulations, we directly used the original empirical branch lengths.

**Table 1:** Empirical phylogenies Dataset

| Dataset | $N^1$ | Tree length | Mean branch length (± sd.) | Reference |
| --- | --- | --- | --- | --- |
| amine<sup>2</sup> | 3039 | 440.861 | 0.073±0.149 | Spielman *et al.* (2015) |
| camelid<sup>3</sup> | 212 | 15.428 | 0.037±0.049 | Kosakovsky Pond and Muse (2005); Murrell *et al.* (2012b, 2013) |
| vertrho<sup>4</sup> | 38 | 12.815 | 0.183±0.159 | Murrell *et al.* (2012b, 2013) |
| h3<sup>5</sup> | 3854 | 5.951 | 0.0005±0.001 | Meyer and Wilke (2015b) |
| hivrt<sup>6</sup> | 383 | 5.642 | 0.007±0.01 | Murrell *et al.* (2012b) |

### *dN/dS* inference

For each simulated codon frequency distribution, we computed *dN/dS* according to the method outlined in Spielman and Wilke (2015b). For each simulated alignment, we inferred site-specific *dN/dS* values with the HyPhy software (Kosakovsky Pond *et al.* 2005) using several approaches: fixedeffects likelihood (FEL) (Kosakovsky Pond and Frost 2005), FUBAR (Murrell *et al.* 2013), and single ancestral counting (SLAC) (Kosakovsky Pond and Frost 2005). We specified the MG94xHKY85 (Muse and Gaut 1994; Kosakovsky Pond and Frost 2005) rate matrix with F1x4 codon frequencies, which has been shown to reduce bias in *dN/dS* estimation (Spielman and Wilke 2015b). We provide customized HyPhy batchfiles, which enforce the F1x4 codon frequency specification, in the Github repository: https://github.com/sjspielman/dnds_1rate_2rate.

For both FEL and FUBAR, we inferred *dN/dS* with both a one-rate model, in which *dN/dS* is represented by a single parameter, and a two-rate model, in which *dN* and *dS* are modeled by separate parameters (Kosakovsky Pond and Frost 2005). For the one-rate FUBAR inferences, we specified 100 grid points to account for the reduced grid dimensionality caused by ignoring *dS* variation, and we specified the default 20x20 grid for two-rate FUBAR inferences (Murrell *et al.* 2013). We left all other settings as their default values. Similarly, for SLAC inference, we calculated *dN/dS* in two ways. As SLAC enumerates *dN* and *dS* on a site-specific basis, there exist two ways to calculate site-wise *dN/dS* :*dS* can be considered site-specific, or *dS* values can be globally averaged, and this mean can be used to normalize all site-specific *dN* estimates. The former calculations effectively correspond to a two-rate model (SLAC2), and the latter calculations correspond to a one-rate model (SLAC1). We conducted all inferences using the true tree along which we simulated each alignment.

As in Kosakovsky Pond and Frost (2005), we excluded all unreliable *dN/dS* inferences when correlating inferred and true *dN/dS* values. Specifically, we excluded FEL estimates where *dN/dS* = 1 and the *P*-value indicating whether the estimate differed significantly from 1 was also equal to 1. Such estimates represent uninformative sites where no mutation has occured (Meyer *et al.* 2015). In addition, we excluded SLAC2 estimates if the number of synonymous mutations counted was 0, and hence the resulting *dN/dS* was undefined. Finally, we excluded all FEL and FUBAR inferences for which the algorithm did not converge as uninformative.

### Statistical Analysis

Statistics were conducted in the R statistical programming language. Linear modeling was conducted using the R package lme4 (Bates *et al.* 2012). We inferred effect magnitudes and significance, which we corrected for multiple testing, using the `glht()` function in the R package multcomp (Hothorn *et al.* 2008). In particular, we built mixed-effects linear models in the lme4 package with the general code lmer(X *∼* method + (1|replicate) + (1|N:B)), where *X* is either the Pearson correlation between inferred and true *dN/dS* or the root-mean-square deviation (RMSD) of the inferred from the true *dN/dS*. RMSD is calculated as 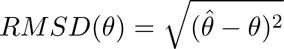 where 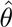 is the true parameter value and *θ* is the estimated value. Note that, for all mixed-effects linear models, we excluded simulations under the *B* = 0.0025 branch length condition.

### Data Availability

All code and results are freely available from the Github repository: https://github.com/sjspielman/dnds_1rate_2rate.

## Results

### Approach

We simulated fully heterogeneous alignments under the HB98 MutSel model (Halpern and Bruno 1998) using the simulation software Pyvolve (Spielman and Wilke 2015a). Our simulation strategy is described in detail in *Methods and Materials*. Briefly, MutSel models are parameterized with a nucleotide-level mutation model and a distribution of codon fitness values. All simulations employed the HKY85 mutation model (Hasegawa *et al.* 1985) with the transition-transversion bias parameter κ= 4.0. We simulated data under four primary conditions: specifying either equal or unequal nucleotide frequencies in the HKY85 model, and specifying no codon bias or codon bias for the codon fitness values. Simulations without codon bias assumed that all synonymous codons had the same fitness, and simulations with codon bias assumed that synonymous codons differed in fitness values. We refer to simulations with equal nucleotide frequencies as ∏_equal_ and to simulations with unequal nucleotide frequencies as ∏_unequal_.

Each simulated alignment contained 100 sites, and simulations were conducted along balanced phylogenies with the number of sequences *N* set as either 128, 256, 512, 1024, or 2048 and with branch lengths *B* set as either 0.0025, 0.01, 0.04, 0.16, or 0.64. For each of the 25 possible combinations of parameters *N* and *B*, we simulated 50 replicate alignments. Importantly, the site-specific evolutionary models were the same within each simulation set, making inferences across *N* and *B* conditions directly comparable. We note that the extremely high divergence level in *B* = 0.64 simulations does not represent real sequence data, but these simulations do allow us to study *dN/dS* inferences under the limiting condition of (approximately) infinite time.

We inferred site-specific *dN/dS* for each simulated alignment using three approaches: fixedeffects likelihood (FEL) (Kosakovsky Pond and Frost 2005), single-likelihood ancestor counting (SLAC) (Kosakovsky Pond and Frost 2005), and FUBAR (Murrell *et al.* 2013). Each of these methods employs a somewhat different approach when computing site-specific *dN/dS* values. FEL fits a unique *dN/dS* model to each alignment site (Kosakovsky Pond and Frost 2005), SLAC directly counts nonsynonymous and synonymous changes along the phylogeny where ancestral states are inferred with maximum likelihood (Kosakovsky Pond and Frost 2005), and FUBAR employs a Bayesian approach to determine *dN/dS* ratios according to a pre-specified grid of rates (Murrell *et al.* 2013).

For each inference method, we inferred *dN/dS* at each site in both a two-rate context (separate *dN* and *dS* parameters per site) and in a one-rate context (a single *dN/dS* parameter per site). Although SLAC, as a counting-based method, always enumerates both *dN* and *dS* on a per-site basis, one can derive an effectively one-rate SLAC by normalizing each site-wise *dN* estimate by the mean of all site-wise *dS* estimates. We refer to one-rate inferences with these methods as FEL1, FUBAR1, and SLAC1, and similarly to two-rate inferences as FEL2, FUBAR2, and SLAC2, respectively. Throughout, we use *method* to refer to distinct algorithmic approaches (FEL, FUBAR, and SLAC), and we use *model* to refer to a one-rate or a two-rate parameterization. We use either *framework* or *approach* to more generally discuss one-rate vs. two rate methods.

We performed all *dN/dS* inference using the HyPhy batch language (Kosakovsky Pond *et al.* 2005). Note that we did not consider the popular random-effects likelihood methods introduced by Yang *et al.* (2000) (e.g. M3, M5, M8) because these methods are used predominantly in a one-rate context. Available two-rate extensions to this framework are computationally burdensome and cannot model the amount of rate heterogeneity required to calculate per-site rates (Kosakovsky Pond and Muse 2005). Finally, we computed true *dN/dS* values from the MutSel parameters, using the approach described in Spielman and Wilke (2015b).

### Modeling synonymous rate variation may reduce inference accuracy

After inferring site-wise *dN/dS* for all simulated alignments, we quantified performance for all inference frameworks using several metrics, notably the Pearson correlation between true and inferred *dN/dS* and the root-mean-square deviation (RMSD) of inferred from true *dN/dS*. Importantly, our simulation strategy necessitates a somewhat different interpretation of results than would more traditional simulation approaches. In particular, the true *dN/dS* ratios calculated from the MutSel parameterizations used during simulation correspond to the *dN/dS* expected at steady state, which in turn indicates the signature of natural selection at evolutionary equilibrium. We can only expect to recover this true *dN/dS* value if the simulated data reflect this condition of stationary. When either the simulated divergence or number of sequences analyzed is low, then, it not necessarily possible to capture the true equilibrium distribution of codons. Therefore, to determine the relative performance of *dN/dS* inference methods, we considered the most accurate inferences as those with the highest *dN/dS* correlations, or conversely the lowest RMSD, within a given choice of *N* and *B*.

In Figure 1, we show resulting Pearson correlation coefficients, averaged across all 50 replicates, between inferred and true *dN/dS* for each inference framework, specifically for ∏_unequal_ simulations. Results for ∏_equal_ simulations showed virtually identical correlations (*P* = 0.68 ANOVA, Figure S1). In the absence of codon bias, *dS* was equal to 1 at all sites. As such, we expected that one-rate inference frameworks would outperform two-rate inference frameworks. We indeed found that one-rate inference frameworks showed the highest correlations when there was no synonymous selection (Figure 1A), in particular at low-to-intermediate divergence levels (*B* ∈ {0.01, 0.04, 0.16}). As the sequences became more diverged, and hence more informative, two-rate frameworks increasingly performed as well as one-rate frameworks did. Even so, two-rate frameworks almost never outperformed one-rate frameworks.

**Figure 1:**
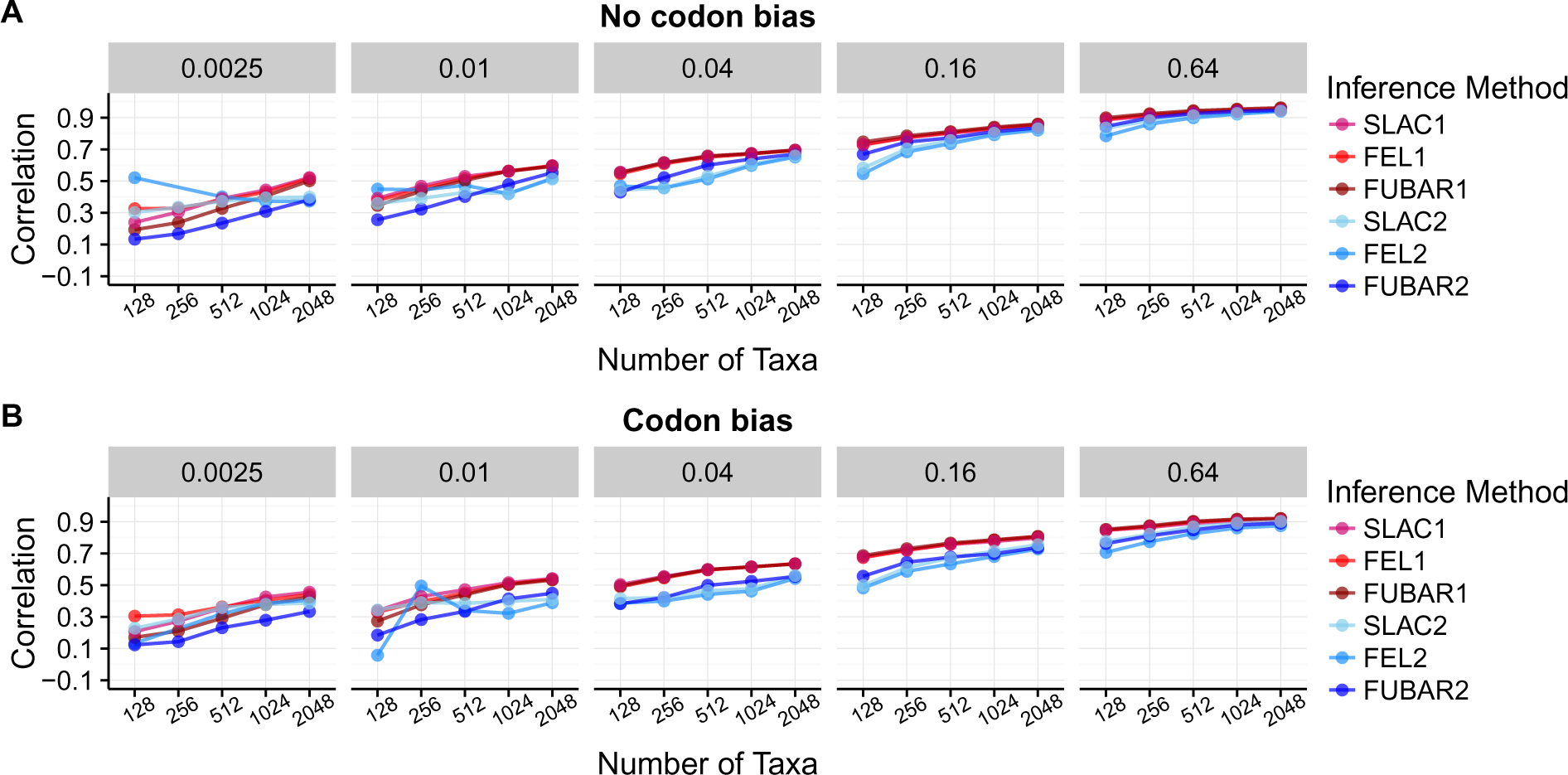
Pearson correlation coefficients between true and inferred *dN/dS* across inference approaches and *N*-*B* conditions, for ∏_unequal_ simulations. A) Correlations for alignments simulated without codon bias. B) Correlations for alignments simulated with codon bias. The label above each sub-plot indicates the branch lengths *B* of the balanced phylogeny along which sequences were simulated, and the x-axes indicate the number of sequences *N*. Each point represents the correlation coefficient averaged across 50 replicates. A corresponding figure for ∏_equal_ simulations is in Figure S1. Note that certain FEL2 points (at *B* = 0.0025 and *B* = 0.01/*N* = 128, for codon bias simulations) are not present because FEL2 generally failed to converge under these conditions.

In the presence of codon bias, both *dN* and *dS* varied at each site, and therefore we expected that two-rate frameworks would be more well-suited for these simulations. Surprisingly, however, one-rate frameworks still outperformed two-rate frameworks across *N* and *B* conditions, in spite of the pervasive site-wise *dS* variation across sites (Figure 1B and Figure S1). Moreover, correlation differences between one-rate and two-rate frameworks were more pronounced for simulations with codon bias than for simulations without codon bias. In other words, two-rate frameworks performed worse on data simulated with codon bias than they did on data simulated without codon bias.

To complement our correlation analysis, we calculated several additional metrics to quantify accuracy: i) root-mean-square deviation (RMSD) of the inferred *dN/dS* from the true *dN/dS* (Figure S2), ii) estimator bias for each inference framework (Figure S3), and iii) variance in residuals for a simple linear model regressing inferred on true *dN/dS* (Figure S4). These metrics displayed the same general trends as did correlation analysis: One-rate frameworks were generally more accurate and precise (lower RMSD, lower estimator bias, and lower residual variance) than were two-rate frameworks, and these overarching trends were more pronounced for simulations with codon bias (all *P* < 2 × 10^*−*16^, ANOVA). As divergence increased, each metric dropped substantially for both one- and two-rate frameworks, with error and/or bias for one-rate frameworks dissipating more quickly than for two-rate frameworks. These patterns were consistent between the ∏_unequal_ and ∏_unequal_ simulations for estimator bias and residual variance (both *P* > 0.2, ANOVA), although ∏_equal_ displayed marginally smaller RMSD values compared to ∏_unequal_ (*P* = 0.04 with an average difference of − 4.45 × 10^−3^, ANOVA). Thus *dN/dS* inference was robust to the presence of nucleotide compositional bias.

### Rate parameterization affects *dN/dS* estimates more strongly than does inference method

We next quantified performance differences among inference frameworks more rigorously, using linear models. For each simulation set, we built mixed-effects linear models with either Pearson correlation or RMSD as the response, inference approach as a fixed effect, and replicate as well as interaction between *N* and *B* as random effects. We performed multiple comparisons tests, with corrected *P*-values, to ascertain the relative performance across inference approaches.

Linear model analysis confirmed prior observations that each one-rate method significantly outperformed its respective two-rate counterpart (Figure 2 for ∏_unequal_ simulations, and Figure S5 for ∏_equal_ simulations). In addition, correlations differences among one-rate methods were not statistically significant for any inferences performed on ∏_unequal_ simulations (Figure 2A). For ∏_equal_ simulations, SLAC1 yielded significantly higher correlations than did FUBAR1, although the effect magnitude size was minimal, with a mean difference of *r* = 0.01 (Figure S5A).

**Figure 2:**
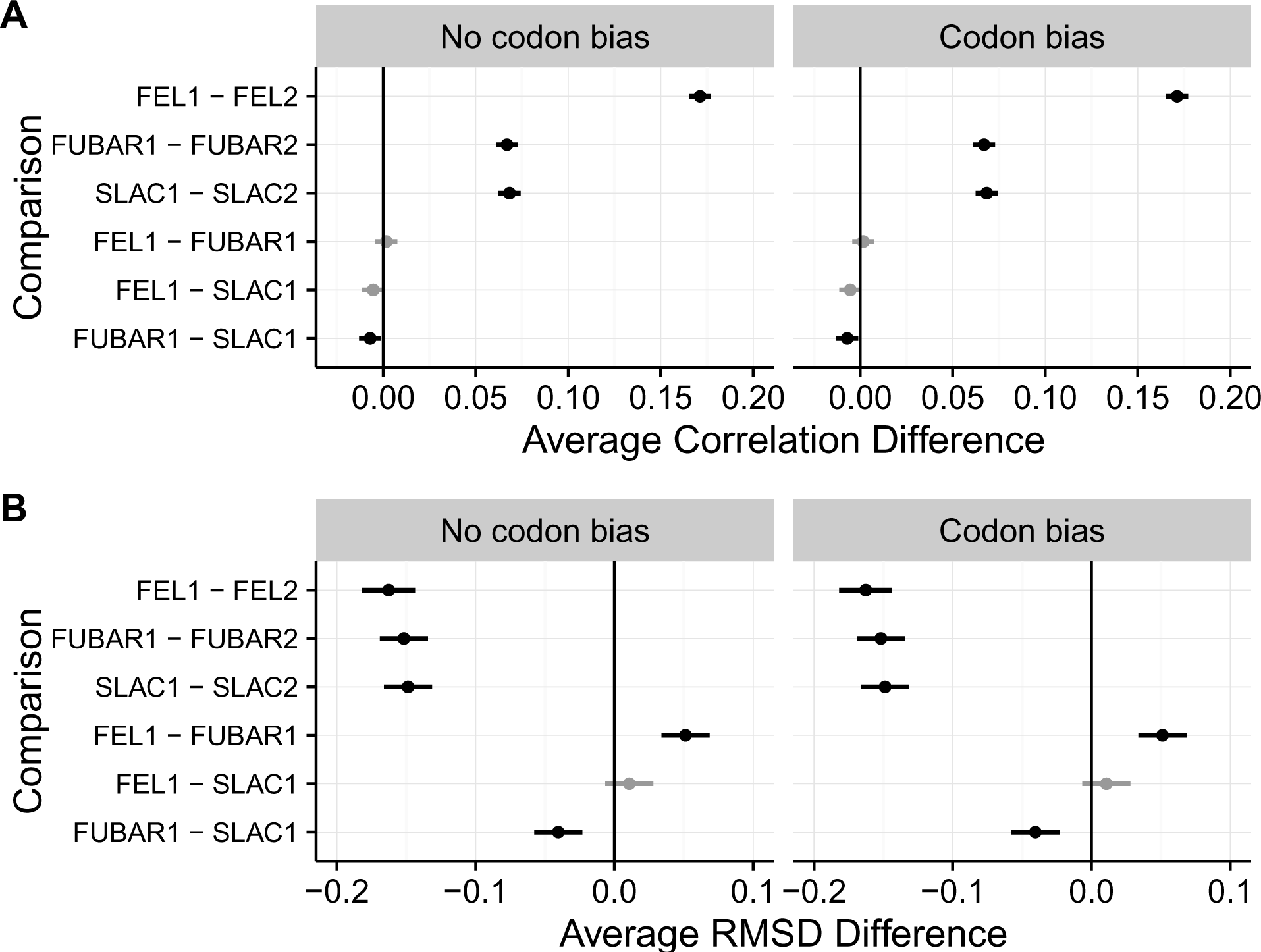
Pairwise comparisons of correlation strength, RMSD, and variance in residual across methods, as determined through multiple comparisons tests, for ∏_unequal_ simulations. A) Results for multiple comparison tests of correlation strength. B) Results for multiple comparison tests of RMSD. Points indicate the estimated average difference between measurement for the respective inference approaches, and lines indicate 95% confidence intervals. Black lines indicate that the performance difference between methods differed significantly from 0 (all *P* < 0.01). Gray lines indicate that the difference was not statically significant (*P* > 0.01). All *P*-values were corrected for multiple testing. Corresponding multiple comparison results for ∏_equal_ simulations are in Figure S5.

Results for RMSD linear models additionally showed, for both ∏_unequal_ and ∏_equal_ simulations, that one-rate methods yielded less error in point estimates than did two-rate methods (Figure 2B and S5B). Unlike results from linear models with correlation as the response, however, RMSD analysis showed some more substantial differences among one-rate methods. Overall, SLAC1 and FEL1 performed comparably, but FUBAR1 showed lower RMSD than both SLAC1 and FEL1, albeit with a very small effect magnitude. This result persisted across all simulation conditions. Together, these findings suggest the number of parameters used to model *dN/dS* mattered more than did the specific inference method chosen.

We next examined whether linear modeling results, specifically those comparing correlation strength between methods were driven by particular simulation conditions. We directly compared correlations between one-rate and two-rate inferences, for each method (FEL, SLAC, and FUBAR) across *N* and *B* conditions, specifically for ∏_unequal_ simulations. These comparisons indicated that improvement of one-rate over two-rate parameterizations was largely driven by results for intermediate divergence levels (Figure S6). For example, under FEL inference, the greatest improvement of FEL1 over FEL2 occurred where *B* ∈ 0.04, 0.16 and *N* ∈ 128, 256, 512.

### Data contain insufficient information for precise site-wise *dS* estimation

We next sought to determine why one-rate frameworks outperformed two-rate frameworks. Given the broad similarly among inference methods and simulation sets, we considered only FEL inferences on ∏_unequal_ simulations for these analyses.

To begin, we confirmed that simulations with codon bias indeed led to *dS* variation in the data. Specifically, we compared distributions of inferred *dS*, with FEL2, between simulations with and without codon bias. If our implementation of codon bias indeed produced *dS* variation, the inferred *dS* distributions from codon bias simulations should contain more variation compared to the inferred *dS* distributions for simulations without codon bias, whose inferences should be concentrated at *dS* ≈ 1. We indeed found that *dS* was much more variable for simulation with codon bias than without (Figure S7), and hence our codon bias simulations did contain substantial site-wise *dS* variation.

We next compared the optimized branch lengths inferred by HyPhy during rate estimation to those used for simulation. Because branch length parameters influence *dN/dS* estimation, it is possible that biases in these parameters could influence the resulting rate inferences. Importantly, we should not expect branch lengths used for a MutSel simulation to match precisely those optimized under a *dN/dS*-based model, due to differences in model assumptions, although branch lengths should be consistently inferred across simulation conditions. Across simulations, we found no significant difference among distribution of optimized branch lengths, for a given set of simulations using the same branch length *B* (Figure S8). Therefore, differences in branch length optimization did not seem to affect *dN/dS* inferences.

We proceeded to compare directly the inferred *dN/dS* values across simulation conditions, for a single representative replicate (Figure S9A-B). The relationship between one-rate and two-rate *dN/dS* estimates featured considerable noise, across all simulation conditions. To determine the source of this noise, we confirmed that *dN* estimates between one-rate and two-rate models were comparable. We examined individual *dN* estimates between FEL1 and FEL2, again for a single replicate. Aside from low-information conditions (e.g.*B* = 0.0025 and/or *N* = 128), *dN* estimates were virtually identical between FEL1 and FEL2, for both simulations with and without codon bias (Figure S9C-D). This result demonstrated that the added *dS* parameter in two-rate inference methods did not affect *dN* estimation, but rather it contributed substantial noise to the final *dN/dS* parameter.

Finally, we assessed how well FEL2 estimated *dN* relative to *dS*, specifically for simulations with codon bias, in which *dS* variation exists. We found that FEL2 consistently estimated *dN* more precisely than *dS*, as measured using both correlations and RMSD (Figure 3). Although accuracy for both *dN* and *dS* estimation increased as either *B* or *N* increased, *dN* estimates universally displayed higher correlations and lower RMSD than did *dS* estimates. As such, it appeared that *dS* was simply statistically more difficult to estimate than was *dN*.

**Figure 3:**
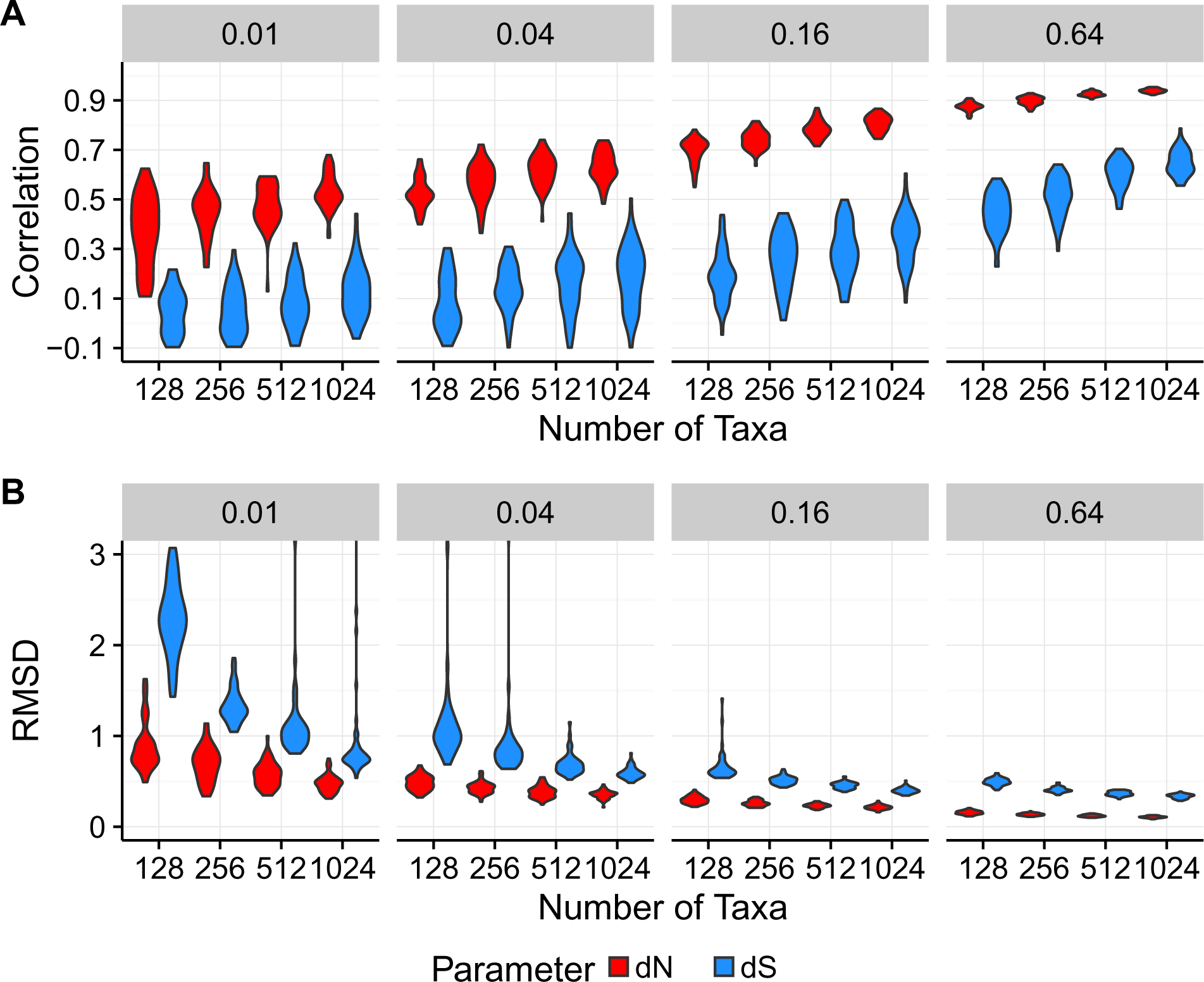
*dN* estimates are more precise than are *dS* estimates. Results in this figure are shown for a subset of conditions for ∏_unequal_ simulations with codon bias, as inferred with FEL2. A) Violin plots of Pearson correlations between inferred and true *dN* and *dS* values. B) Violin plots of RMSD of inferred from true *dN* and *dS* values. Outlying points beyond the y-axis ranges have been removed for visualization.

We hypothesized that this result was a direct consequence of the relative amount of information in the alignments for nonsynonymous vs. synonymous changes. Using the simulated ancestral sequences within each simulated alignment, we directly counted the number of nonsynonymous and synonymous changes which had occurred across the phylogeny. We observed that nonsynonymous changes occurred roughly twice as frequently over the course of a simulation than did synonymous changes (Figure 4). This result was fully compatible with the notion that statistical estimation of *dS* was more challenging than *dN* because of sample size: Alignments contained nearly double the amount of information contributing to *dN* than to *dS*. As a consequence, *dS* estimation was less precise and noisier across simulation conditions, ultimately explaining why two-rate frameworks yielded less precise *dN/dS* estimates compared to one-rate frameworks.

**Figure 4:**
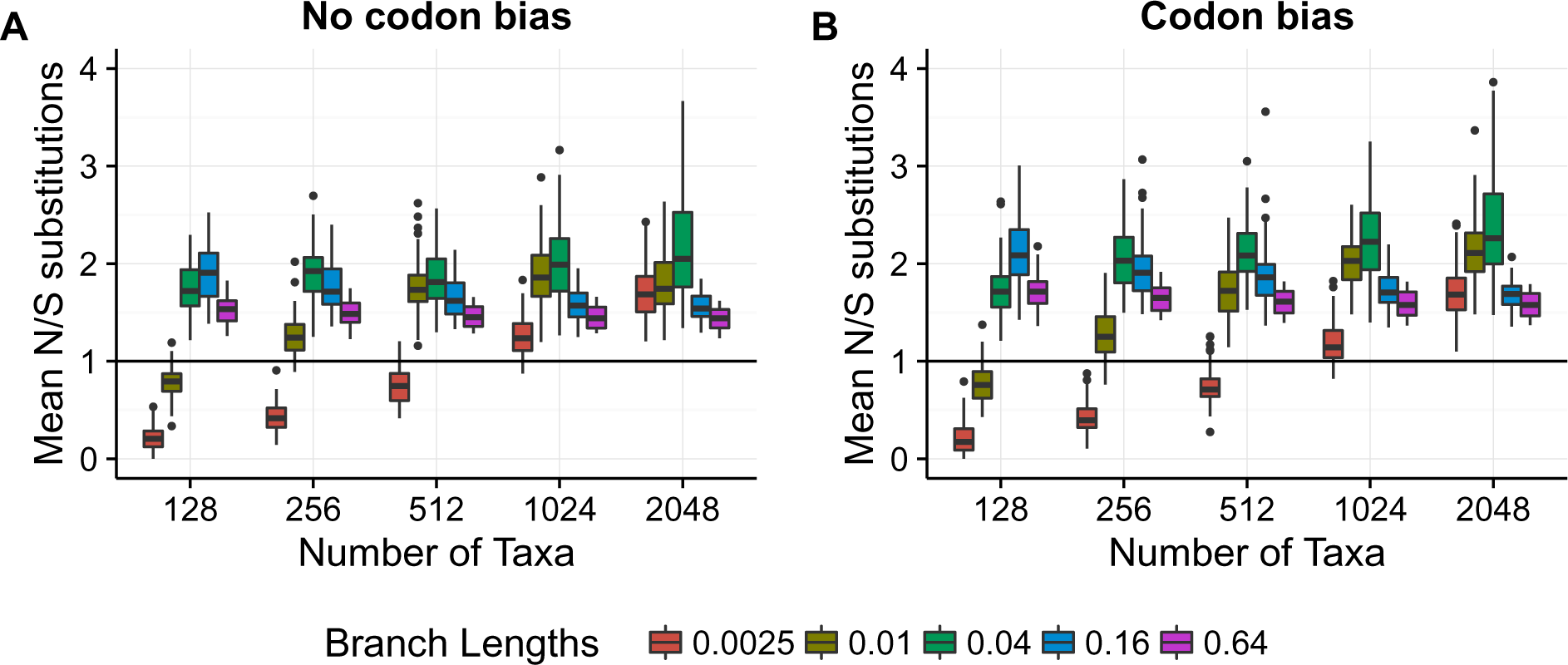
Ratio of the number of nonsynonymous to synonymous changes which occurred during simulations, as counted directly from simulated ancestral sequences. Each boxplot represents, across 50 replicates, the per-alignment ratio of the number of nonsynonymous changes to synonymous changes averaged across sites. Results shown here are from ∏_unequal_ simulations.

### One-rate frameworks outperform two-rate frameworks primarily at weakly-constrained sites

Importantly, all previous analyses compared inferences across alignment replicates. Yet, each alignment featured an array of selective constraints, with each site evolving with a different underlying *dN/dS*. Whether the heterogeneous selective constraints across sites influenced our previous results was not immediately clear. For example, according to the structure of the genetic code, 74% of all possible single-nucleotide changes are nonsynonymous, and the remaining 26% are synonymous. Therefore, a neutrally-evolving site, where both nonsynonymous and synyonymous changes are equally likely to go to fixation, should experience approximately three times more nonsynonyomus than synonymous substitutions. By contrast, sites under stringent selection pressure will tolerate few amino acids, and thus these sites may feature more synonymous than nonsynonymous changes. Noting this distinction, we next examined whether one-rate or two-rate frameworks performed differently depending on a given site’s evolutionary process.

We therefore re-analyzed our ∏_unequal_ simulations, specifically under FEL1 and FEL2 inference, while considering sites to be in one of two categories: Having a relative enrichment for synonymous substitutions or having a relative enrichment for nonsynonymous substitutions (Figure 5). Across simulation conditions, FEL1 and FEL2 models yielded virtually identical correlations at sites enriched for synonymous substitutions (linear model, *P >* 0.4). By contrast, FEL1 consistently outperformed FEL2 when sites contained more nonsynonymous than synonymous substitutions, with an average correlation increase of *r* = 0.1 (linear model, *P* < 2× 10^−16^). The *B* = 0.64 condition did not adhere to this general pattern and consistently favored one-rate frameworks, likely to due difficulty in estimating *dS* due to mutational saturation at such high divergences. Together, these results show that one-rate frameworks offer the most improvement, relative to two-rate frameworks, when sites experience more nonsynonymous changes. On the other hand, when data is enriched for synonymous changes, one-rate and two-rate frameworks provide comparable estimates.

**Figure 5:**
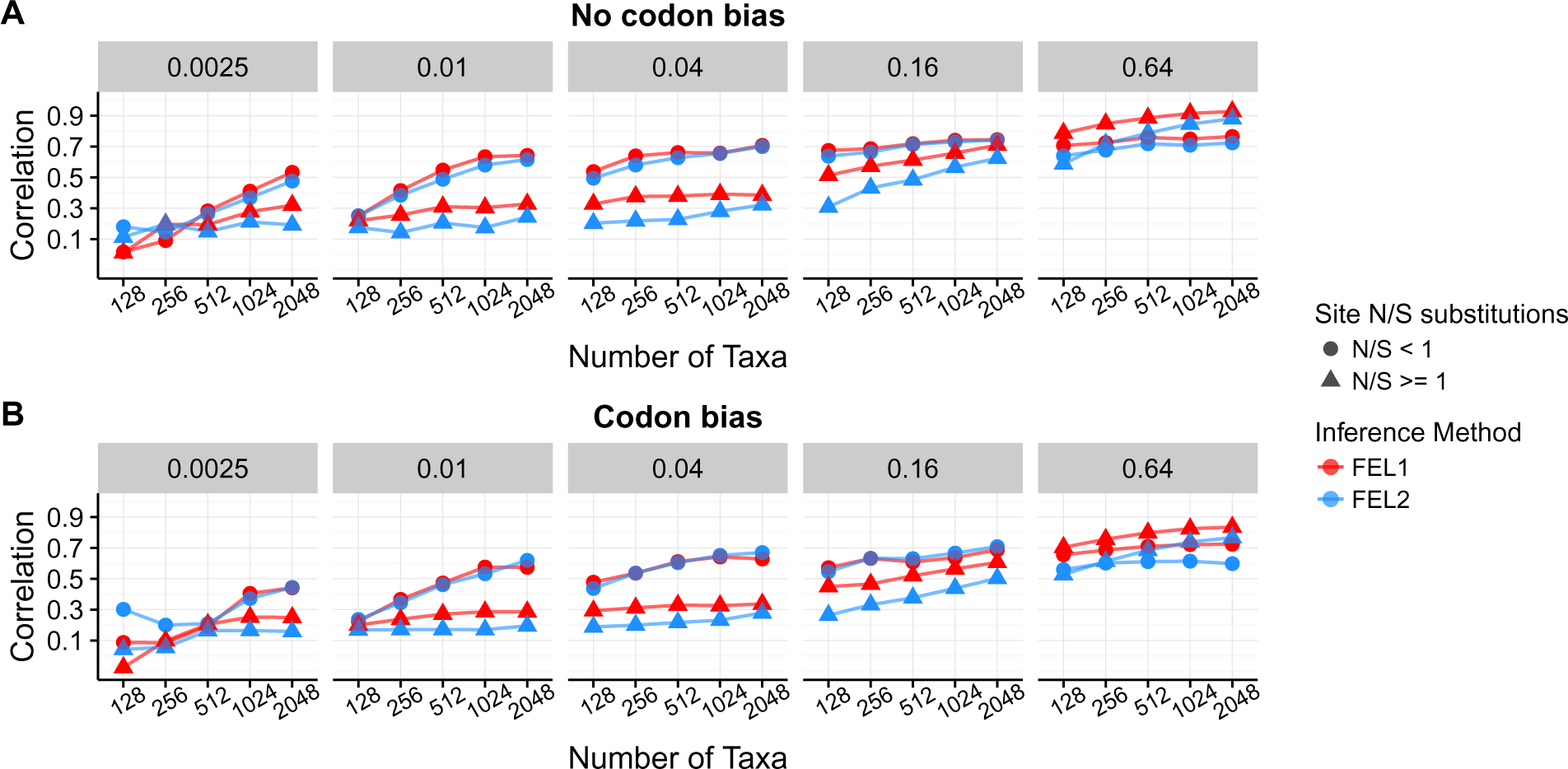
Mean Pearson correlations between inferred and true *dN/dS*, specifically for ∏_unequal_ simulations as analyzed with FEL1 and FEL2. Sites which have experienced relatively more synonymous changes are represented with circles, and sites which have experienced relatively more nonsynonymous changes are represented with triangles.

To ascertain more broadly at which sites a one-rate *dN/dS* inference framework may be preferred, we examined the relationship between substitution counts and true *dN/dS* across simulations (Figure 6). Figure 6A shows the relationship between true *dN/dS* and the mean ratio of nonsynonymous to synonymous substitution counts, along the simulated phylogeny, specifically for *N* = 512 and *B* = 0.04. In these panels, points below the *y* = 1 line represent sites which featured, on average, a relative enrichment for synonymous compared to nonsynonymous changes, and similarly points above the *y* = 1 line represent sites with an average enrichment for nonsynonymous compared to synonymous changes. We found that sites under stronger selective constraint indeed featured relatively more nonsynonymous changes, and sites under weaker constraint featured relatively more synonymous changes.

**Figure 6:**
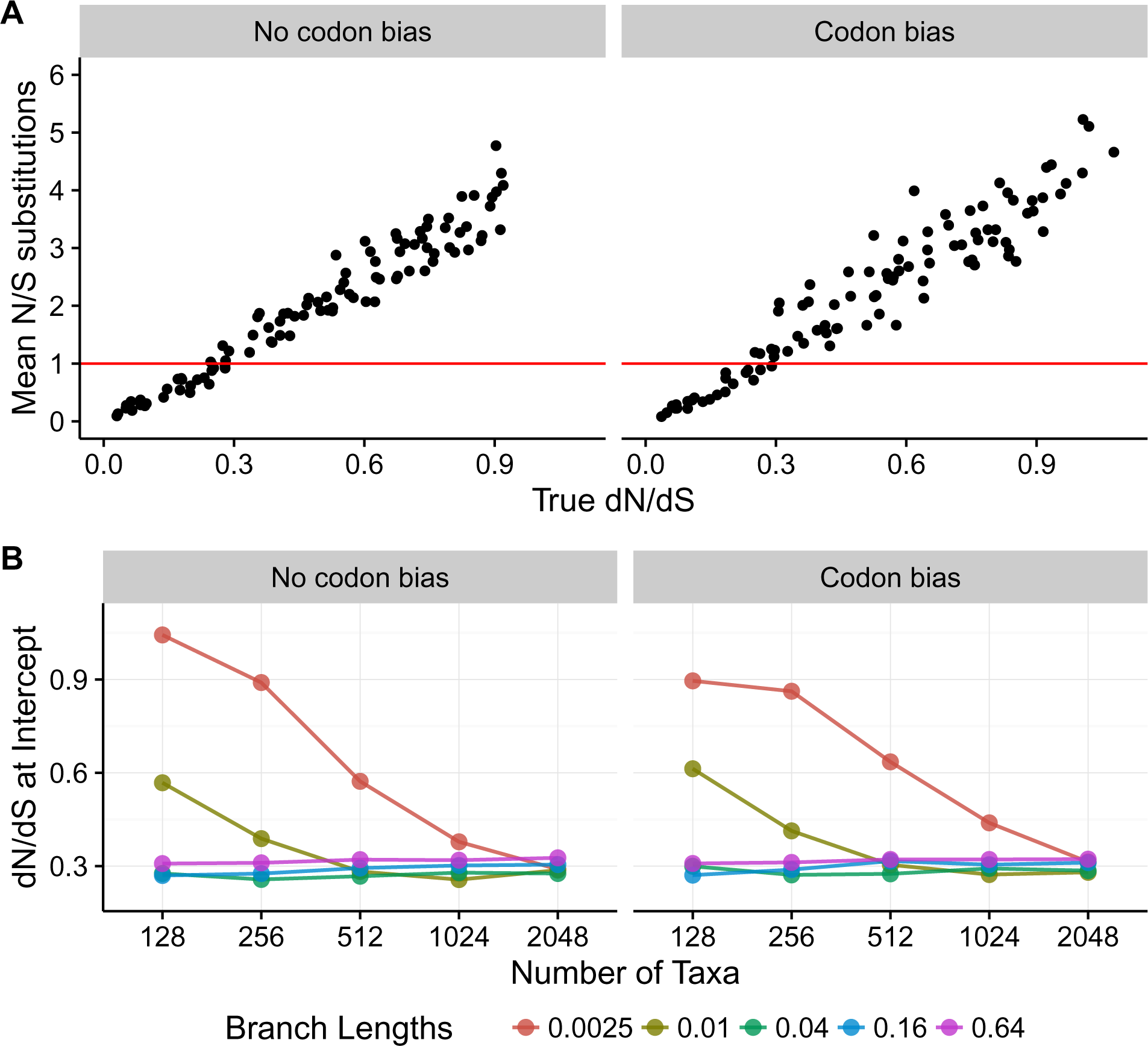
Relationship between true *dN/dS* and the ratio of the number of nonsynonymous to synonymous changes which occurred during simulations, as counted directly from simulated ancestral sequences. Results are shown for ∏_unequal_ simulations. (A) Regression of the ratio of nonsynonymous to synonymous substitution counts specifically for the *B* = 0.04 and *N* = 512 simulation. (B) True *dN/dS*, across simulation conditions, where sites transition from being enriched for synonymous changes to being enriched for nonsynonymous changes.

To generalize across all simulation conditions, we calculated the true *dN/dS* where, on average, sites transitioned from having more synonymous to more nonsynonymous changes (Figure 6B). In general, sites became enriched for nonsynonymous substitutions at *dN/dS* ≈ 0.3. However, the transition point was substantially larger for simulation conditions with low levels of divergence, likely because substitutions did not have sufficient time to accumulate. Taken together, these results reveal that one-rate frameworks may offer the most improvement over two-rate frameworks when *dN/dS* ≥ 0.3, i.e. when sites are under moderate-to-weak purifying selection. By contrast, one-rate and two-rate frameworks show minimal, if any, differences when applied to sites subject to strong purifying selection (*dN/dS* < 0.3).

### Divergence is more important than is the number of sequences for identifying long-term evolutionary constraint

We additionally observed that *dN/dS* inference accuracy increased both as the number of sequences *N* and the branch lengths *B* (divergence) grew (Figures 1, S1–4), suggesting that large and/or highly informative datasets are necessary for the inferred *dN/dS* to capture the actions of natural selection at evolutionary equilibrium. However, it was not immediately clear whether *N*, *B*, or some combination of these conditions drove this trend. Therefore, we next assessed the relative importance of *N* and *B* on estimating the equilibrium *dN/dS* rate ratio.

We calculated the tree length for each *N* and *B* parameterization, specifically for ∏_unequal_ simulations. Note that, in the context of our mutation–selection simulations, the tree length indicates the expected number of substitutions per site across the entire tree, relative to the number of neutral substitutions (Tamuri *et al.* 2012; Spielman and Wilke 2015a). If *N* and *B* served roughly equal roles in terms of providing information, then any combination of *N* and *B* corresponding to the same tree length should have produced similar *dN/dS* correlations. We did not, however, observe this trend; instead, all else being equal, *B* had a significantly greater influence than did *N* on the resulting correlations. For example, as shown in Figure 7, we compared *dN/dS* correlations and RMSD from FEL1 for three combinations of *N* and *B* conditions that all yielded virtually the same tree lengths (162–164). Simulations with lower *N* and higher *B* resulted in far more accurate *dN/dS* estimates, even though all simulations in Figure 7 experienced the same average number of substitutions. This increase was highly significant; for data simulated without codon bias, correlations increased an average of ∼28%, from *B* = 0.04 to *B* = 0.64, and similarly RMSD decreased an average of ∼50% (both *P* < 10^−15^, linear model). For data simulated with codon bias, correlations increased an average of ∼33%, from *B* = 0.04 to *B* = 0.64, and RMSD decreased an average of ∼52% (both *P <* 10^−15^, linear model). Therefore, the relative importance of divergence over number of taxa held for simulations with and without codon bias alike.

**Figure 7:**
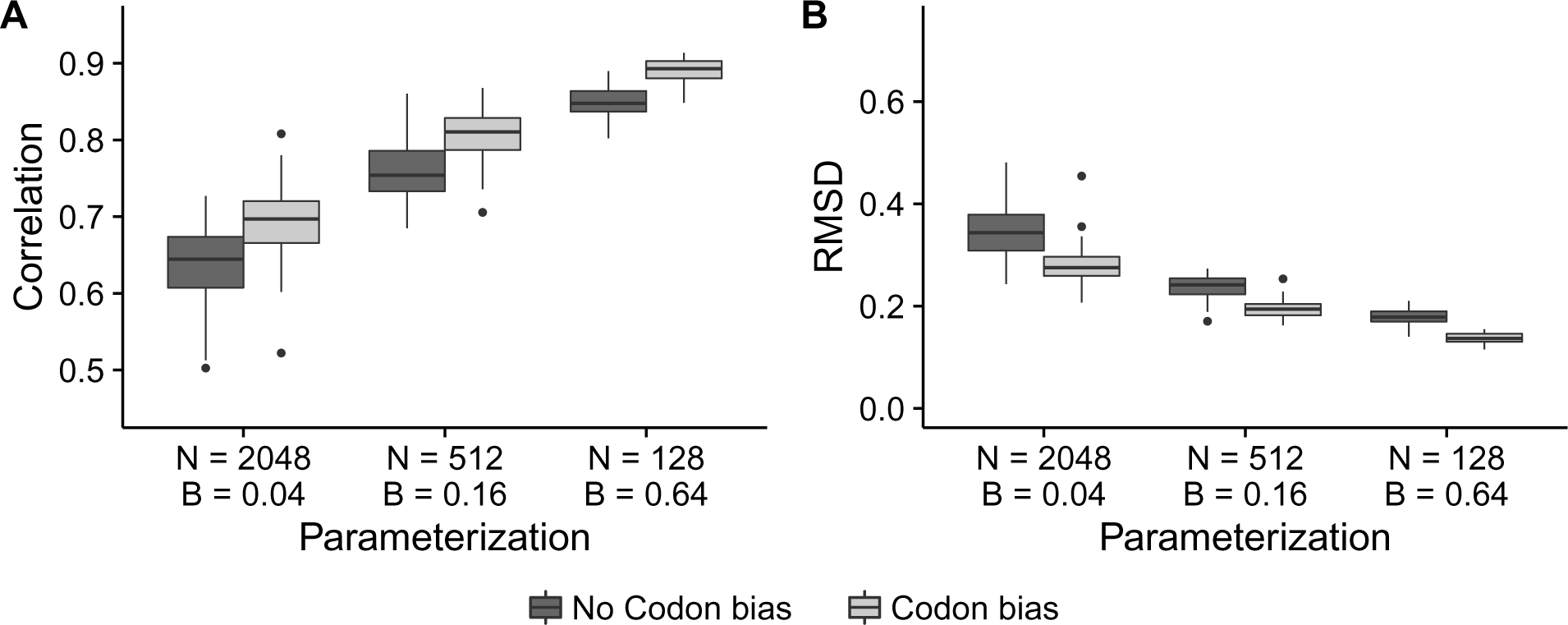
The amount of divergence is more important than the number of sequences is for obtaining the equilibrium *dN/dS* value. Boxplots represent either A) correlation or B) RMSD across the 50 respective simulation replicates. Results shown correspond to ∏_unequal_ simulations as inferred with FEL1. From left to right, tree lengths are equal to 163.76, 163.52, and 162.56. A corresponding figure for ∏_equal_ simulations is in Figure S10.

### Simulations along empirical phylogenies recapitulate observed trends

While the balanced-tree simulations described above provided a useful framework for examining broad patterns in inference-framework behaviors, they did not necessarily reflect the properties of empirical datasets. We therefore assessed how applicable our results were to real data analysis by simulating an additional set of alignments along five empirical phylogenies (Table 1). Importantly, we considered the original empirical branch lengths for this analysis, so these phylogenies featured a range of number of taxa and divergence levels representative of empirical studies. We simulated alignments using the ∏_unequal_ MutSel parameterizations, with both no codon bias and codon bias. We simulated 50 replicate alignments for each phylogeny and parameterization, and for simplicity we inferred *dN/dS* using only FUBAR1 and FUBAR2.

We identified the same general trends in these empirical simulations as we observed for simulations along balanced trees: FUBAR1 estimated *dN/dS* more precisely than did FUBAR2, and phylogenies with higher divergence levels (e.g. longer branch lengths) yielded more accurate estimates (Figure 8). Furthermore, FUBAR2 estimated *dN* more accurately than *dS* (Figure S11), again reflecting the relative difficulty in estimating *dS* compared to *dN*. Importantly, most empirical phylogenies showed mean correlations with true *dN/dS* of 0.4< *r* < 0.6, with the key exception of the biogenic amine receptor phylogeny (”amine”), whose extremely high number of taxa and divergence yield exceptionally high correlations with both FUBAR1 and FUBAR2. Therefore, we found that under more realistic conditions, estimated *dN/dS* will correlate with the equilibrium *dN/dS* ratio with only moderate strength.

**Figure 8:**
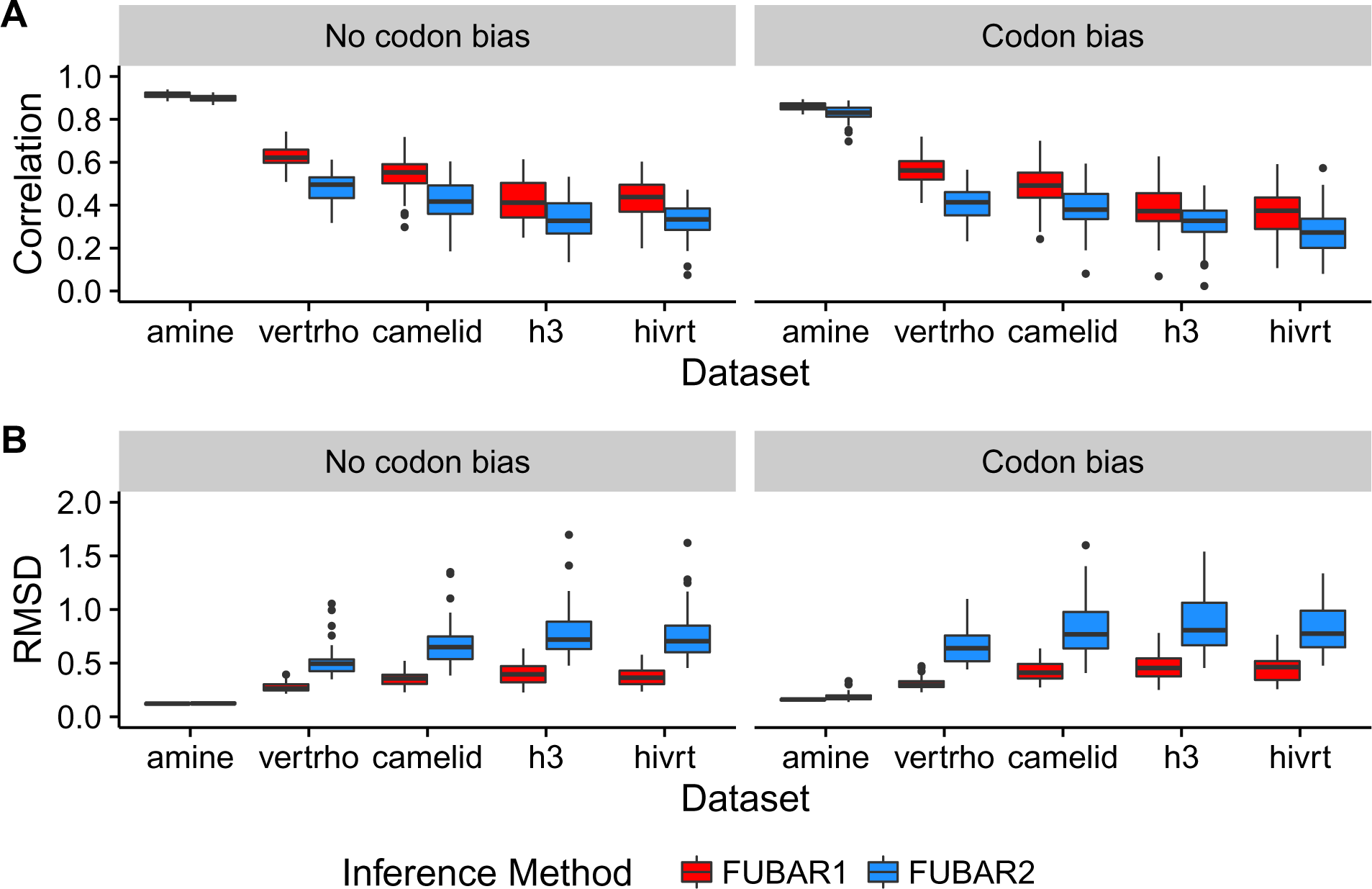
Inference results for simulations performed along empirical phylogenies (Table 1). A) Pearson correlations between inferred and true *dN/dS* values. B) RMSD of inferred from true *dN/dS* values.

In addition, results for simulations along the empirical phylogenies supported our findings regarding the relative importance of divergence vs. number of taxa, in particular through the juxtaposition of results for camelid and vertebrate rhodopsin (”vertrho”) simulations. These phylogenies showed similar tree lengths, but the camelid tree length was driven by the number of sequences, and the vertebrate rhodopsin tree length was driven by its larger branch lengths (Table 1). Correlations and RMSD revealed a far higher inference accuracy for the vertebrate rhodopsin simulations than for the camelid simulations. On average, vertebrate rhodopsin correlations were 0.08 higher than were camelid correlations, and vertebrate rhodopsin RMSD was 0.1 lower than was camelid RMSD (both *P* < 5 × 10^−11^, linear model). These metrics were consistent between simulations with and without codon bias (both *P* > 0.5, linear model test for interaction effect of codon bias parameterization). Therefore, even for data simulated along empirical phylogenies, sequence divergence proved more important than did the number of taxa for accurately estimating the equilibrium *dN/dS* rate ratio.

## Discussion

In this study, we have examined the relative accuracy of one-rate and two-rate site-specific *dN/dS* inference approaches. Importantly, we performed these analyses in the specific context of *dN/dS* point estimation in the presence of model misspecification. We have found that one-rate inference models usually yielded more accurate inferences than do two-rate models, across a variety of inference algorithms. More specifically, we provide evidence that one-rate models may improve upon two-rate models predominantly for sites subject to moderate-to-weak purifying selection. By contrast, one-rate and two-rate models infer *dN/dS* point estimates with comparable accuracy when sites are under stronger purifying selection. These results hold for both sequences without codon bias (synonymous codons are equally fit) and with codon bias (synonymous codons differ in fitness), suggesting that two-rate models are not necessarily more reliable than are one-rate models even when *dS* variation exists. We attribute these results to the relative amounts of information present in the data used for estimating *dN* and *dS* parameters. When relatively more information is available to estimate *dN*, the *dS* parameter becomes overly-influenced by noise and hence reduces accuracy in *dN/dS* estimates.

Our study provides novel insight into how *dN/dS* inference frameworks behave specifically when they are misspecified to the data. Indeed, real genomes do not evolve according to a *dN/dS* model, and thus virtually all applications of *dN/dS* models will be misspecified to some degree. Although it is certainly true that mutation–selection models also do not precisely capture real sequence evolution, this simulation framework provides a starting place to uncover model properties and limitations in an explicitly misspecified context.

We have demonstrated that two-rate frameworks do not necessarily accomplish their intended goal of modeling synonymous rate variation. Logically, one would presume that, when *dS* differs among sites, estimating *dS* separately across sites would produce more accurate *dN/dS* estimates than would fixing *dS* to a constant value. Indeed, an assumed presence of synonymous substitution-rate variation is the very justification for using a two-rate *dN/dS* model (Kosakovsky Pond and Muse 2005). However, if the data do not contain sufficient information for inferring *dS*, then two-rate parameterizations may suffer from excessive amounts of noise, and hence in certain circumstances, one-rate models may be preferable.

We additionally have found that high levels of sequence divergence are critically important for obtaining a reliable steady-state *dN/dS* value, moreso than the number of sequences analyzed (Figure 7). This finding has important implications for data set collection: It may be preferable to include fewer, more divergent sequences rather than as many sequences as one can obtain. More specifically, increasing the number of taxa in a given analysis may only be beneficial if the new sequences are substantially diverged from the existing sequences. Measuring *dN/dS* from thousands of sequences with low divergence may actually be less effective than analyzing fewer, more diverged sequences, even if the mean number of per-site substitutions would be the same.

Our use of MutSel models for simulations raises several important caveats that directly impact how to interpret our results. As described, our results are contingent on the fact that our simulation model did not match our inference model, and hence the inference model was mathematically misspecified. In other words, we ask how *dN/dS*-based models perform in estimating a parameter that the MutSel model does not explicitly contain, although it can be calculated from the MutSel parameters. As a consequence, certain biases arise during *dN/dS* inference. For example, comparable performance of SLAC, an approximate counting-based method, with FEL and FUBAR, both of which employ more rigorous statistical procedures, may be directly attributed to model misspecification. Indeed, previous studies using data simulated under the inference model have suggested that SLAC may be a biased estimator when correctly specified, particularly at high divergences (Kosakovsky Pond and Frost 2005). Therefore, it is certainly possible that a two-rate *dN/dS* framework would outperform a one-rate *dN/dS* framework if the model were correctly specified. Indeed, previous studies have shown that two-rate *dN/dS* models perform well for data simulated with explicit *dN* and *dS* constant parameters (Kosakovsky Pond and Muse 2005; Kosakovsky Pond and Frost 2005; Murrell *et al.* 2012c, 2013).

Second, the substitution process under a MutSel models is not necessarily temporally homogeneous, whereas in *dN/dS*-based models, substitutions are Poisson-distributed over time. As a consequence, sequence divergence will have a strong effect on inference accuracy for data generated in a MutSel framework. Indeed, *dN/dS* can only be calculated based on the substitutions which have occurred in the sequence data examined. If, for example, sequences were not highly diverged, then *dN/dS* estimates will be biased based on which substitutions have had an opportunity to occur. Conversely, for data simulated under a *dN/dS* framework, in which all nonsynonymous substitutions occur at the same rate, it will matter far less which substitutions had a chance to occur, as all proceed at the same rate. Therefore, because the temporal inhomogeneity of the MutSel process does not match the corresponding homogeneity of the *dN/dS*-based model process, our observed correlations between inferred and true *dN/dS* (Figure 1) were generally lower than they would have been if sequences were simulated with a *dN/dS*-based model.

Third, because MutSel models can only correspond to sites evolving under an evolutionary equilibrium [i.e. under either purifying selection or neutral evolution where *dN/dS* ≤ 1 (Spielman and Wilke 2015b)], our results do not necessarily apply to contexts where sequences do not evolve under equilibrium, or for the specific application of positive-selection inference (*dN/dS >* 1). For example, *dN/dS* inference is perhaps most commonly used to study sequences evolving along a changing fitness landscape, as would be the case for viral and/or pathogen evolution (Delport *et al.* 2008; Murrell *et al.* 2012a; Demogines *et al.* 2013; Meyer *et al.* 2015; Meyer and Wilke 2015a) By contrast, the MutSel model used here assumes that the fitness landscape is static across the phylogeny. Therefore, we must emphasize that our results apply primarily to sequences evolving under equilibrium conditions, and not necessarily to sequences evolving under shifting selection pressures. As such, our results may or may not have any bearing on parameterizations used for positive-selection inference.

Finally, our codon bias simulations assumed that *dS* variation was driven by selection on synonymous codons. In other circumstances, synonymous rate variation might emerge when a given gene contains mutational hotspots, e.g. regions with a strongly elevated nucleotide mutation rate. In such circumstances, it is possible that a two-rate model would outperform a one-rate model if the variation in mutational processes were sufficiently large (Kosakovsky Pond and Frost 2005).

Our results builds on the well-documented time-dependency of the *dN/dS* metric, a phenomenon studied primarily in the context of polymorphic data (Rocha *et al.* 2006; Kryazhimskiy and Plotkin 2008; Wolf *et al.* 2009; Mugal *et al.* 2014; Meyer *et al.* 2015). Our results extend these findings and indicate that this time-dependency is more general and pertains also to circumstances where the data contain only fixed differences. This finding makes intuitive sense: As divergence increases, sites will be more likely to visit the full range of selectively tolerated states, at which time the long-term evolutionary constraints will become apparent. It is therefore likely that most *dN/dS* measurements will be biased by time to some degree, even if all differences are fixed and not polymorphic.

Finally, our study has important implications for research that seeks to relate site-specific *dN/dS* ratios to protein structural properties, such as relative-solvent accessibility or weighted contact number (Echave *et al.* 2016). These metrics reflect the overarching biophysical constraints that influence protein evolutionary trajectories. Studies which have examined the correlations between site-specific *dN/dS*, or conversely amino-acid level evolutionary rates, and such structural quantities have recovered correlations with strengths widely ranging from 0.1–0.8 (Shih and Hwang 2012; Huang *et al.* 2014; Yeh *et al.* 2014b, a; Shahmoradi *et al.* 2014; Meyer and Wilke 2015b, c; Jackson *et al.* 2016). Recent work by Jackson *et al.* (2016) has attributed the source of this wide range of correlations directly to the extent of sequence divergence present in a given dataset, such that more diverged datasets display higher correlations and less diverged datasets display lower correlations. Our findings that increased divergence levels contribute strongly to *dN/dS* inference accuracy are fully consistent with those of Jackson *et al.* (2016). Therefore, we suggest that future work examining the relationship between protein evolutionary rate and structure should focus on datasets with intermediate-to-high divergences, which are most likely to provide meaningful information about long-term evolutionary constraints.

## Acknowledgments

This work was supported in part by NIH grant F31 GM113622-01 to SJS, NIH grant R01 GM088344 to COW, ARO grant W911NF-12-1-0390 to COW, DTRA grant HDTRA1-12-C-0007 to COW, and NSF Cooperative Agreement No. DBI-0939454 (BEACON Center) to COW. Computational resources were provided by the University of Texas at Austin’s Center for Computational Biology and Bioinformatics (CCBB) and the Stampede cluster at the Texas Advanced Computing Center (TACC). We thank Julian Echave for insightful discussion and helpful feedback.

## Supplementary Figures

**Figure S1.**
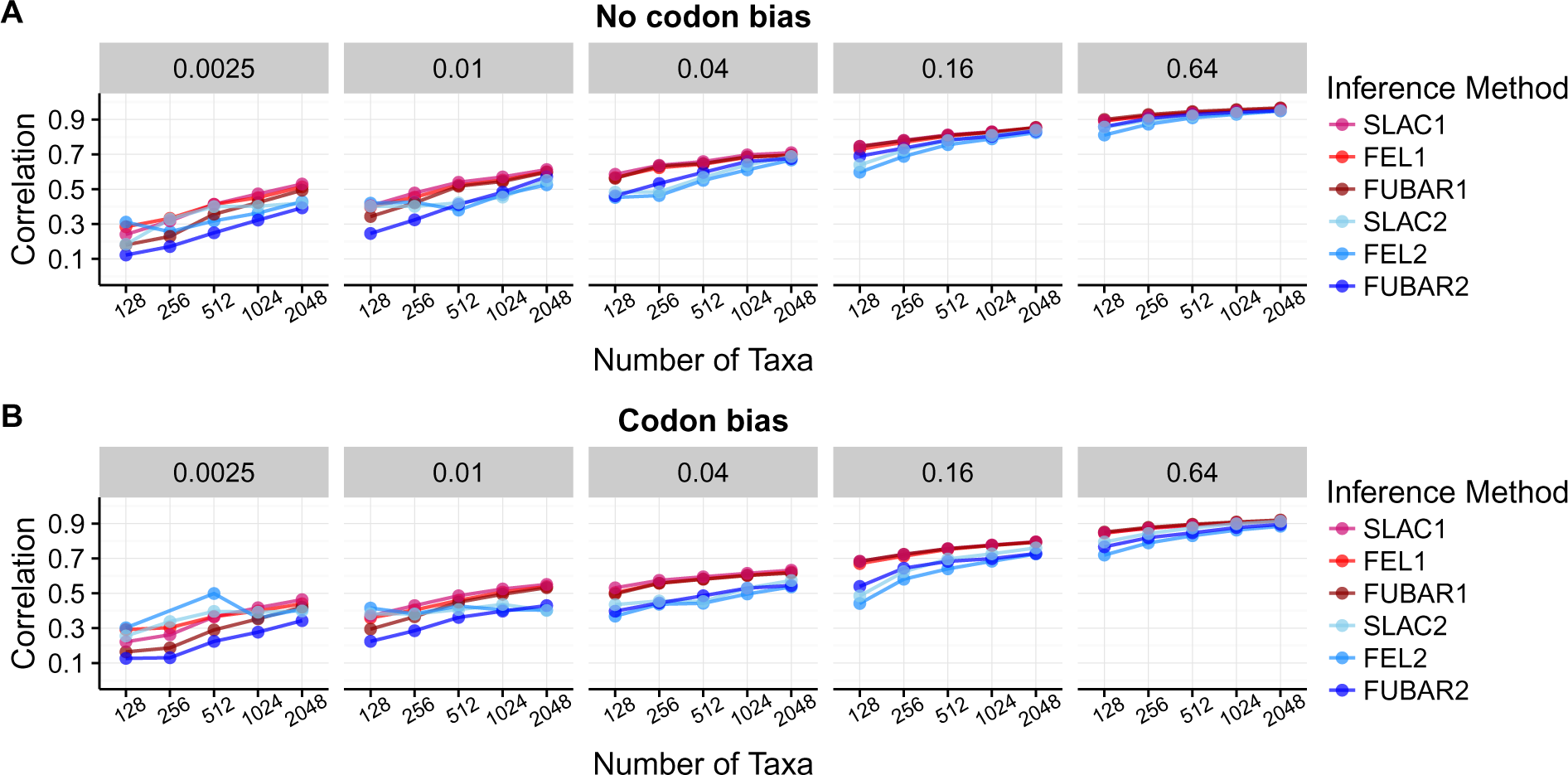
Pearson correlation between true and inferred *dN/dS* across methods and simulation conditions. Data shown in this figure corresponds to ∏_equal_ simulations. A) Correlation between true and inferred *dN/dS* for simulations with no codon bias. B) Correlation between true and inferred *dN/dS* for simulations with codon bias. Note that certain FEL2 points (*B* = 0.0025/ *N 2*{128, 256} and *B* = 0.01/*N* = 128, for codon bias simulations) are not present because FEL2 generally failed to converge under these conditions.

**Figure S2.**
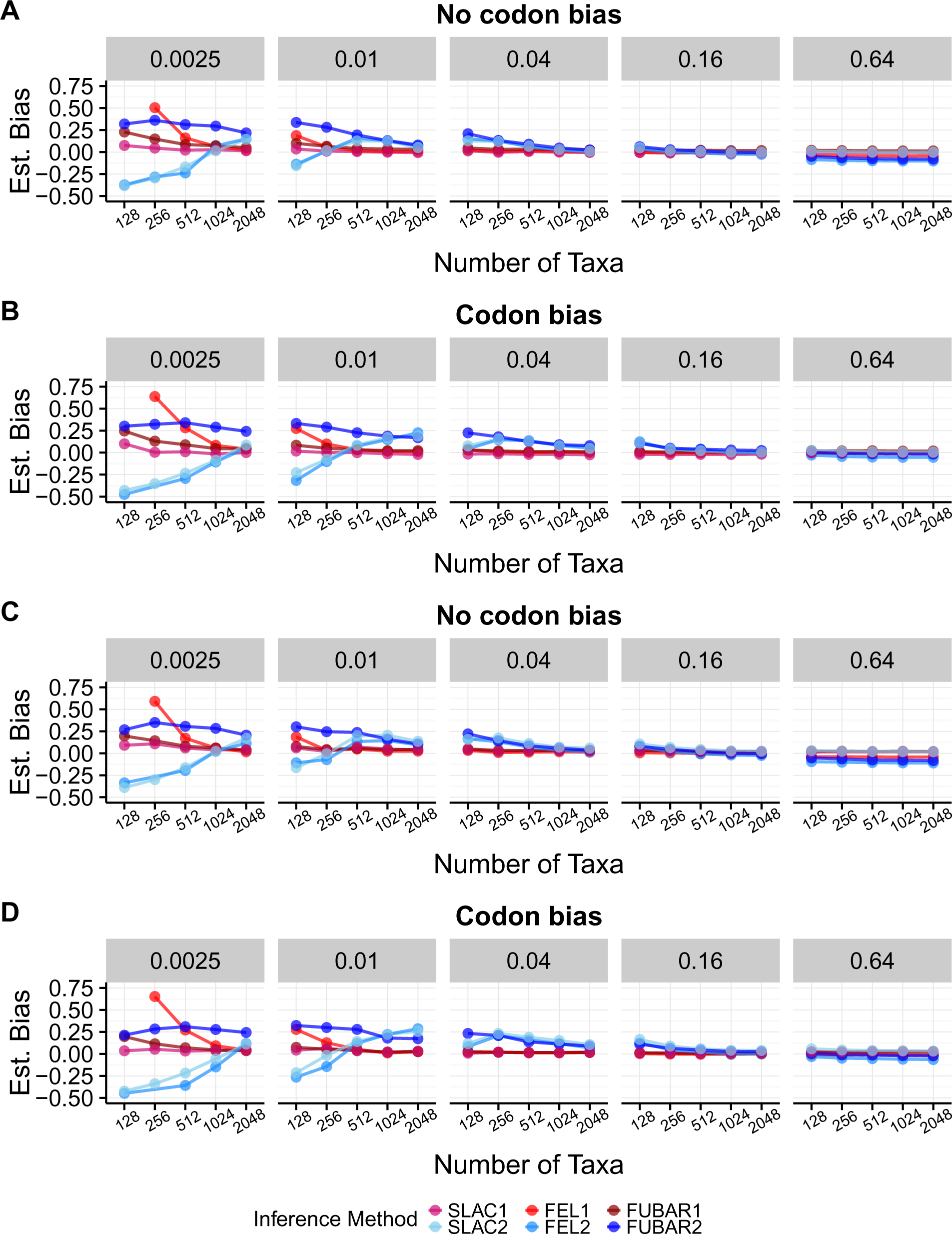
*dN/dS* estimator bias of each inference method inferred across simulation conditions. A) ∏_equal_ simulations with no codon bias. B) ∏_equal_ simulations with codon bias and equal nucleotide frequencies. C) ∏_unequal_ simulations with no codon bias. D) ∏_unequal_ simulations with codon bias. Note that certain FEL1 and FEL2 points are not present in the figure for conditions where the methods generally failed to converge.

**Figure S3.**
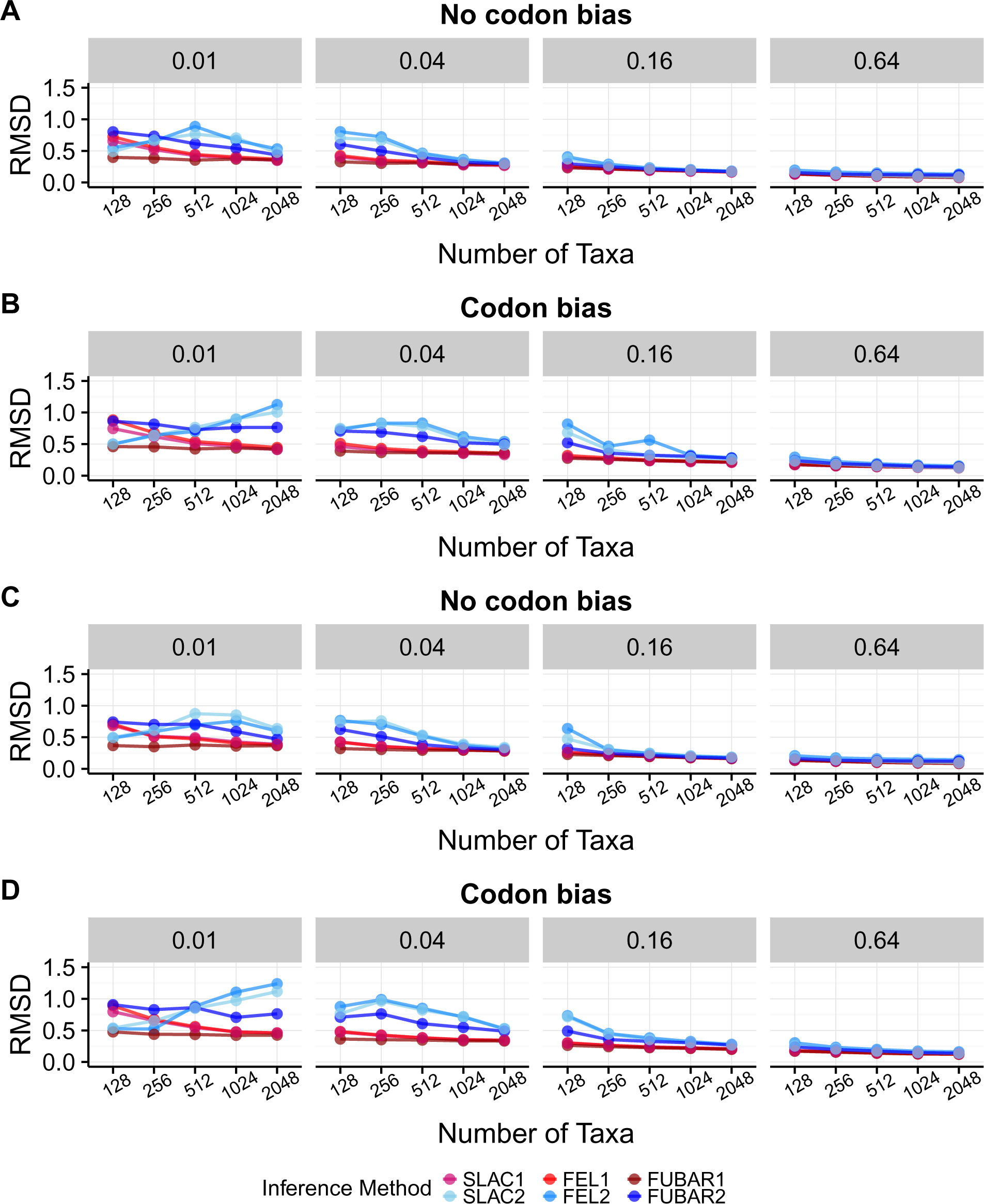
RMSD of inferred from true *dN/dS* across inference methods and simulation conditions. A) ∏_equal_ simulations with no codon bias. B) ∏_equal_ simulations with codon bias and equal nucleotide frequencies. C) ∏_unequal_ simulations with no codon bias. D) ∏_unequal_ simulations with codon bias. Note that certain FEL2 points are not present in the figure for conditions where the methods generally failed to converge.

**Figure S4.**
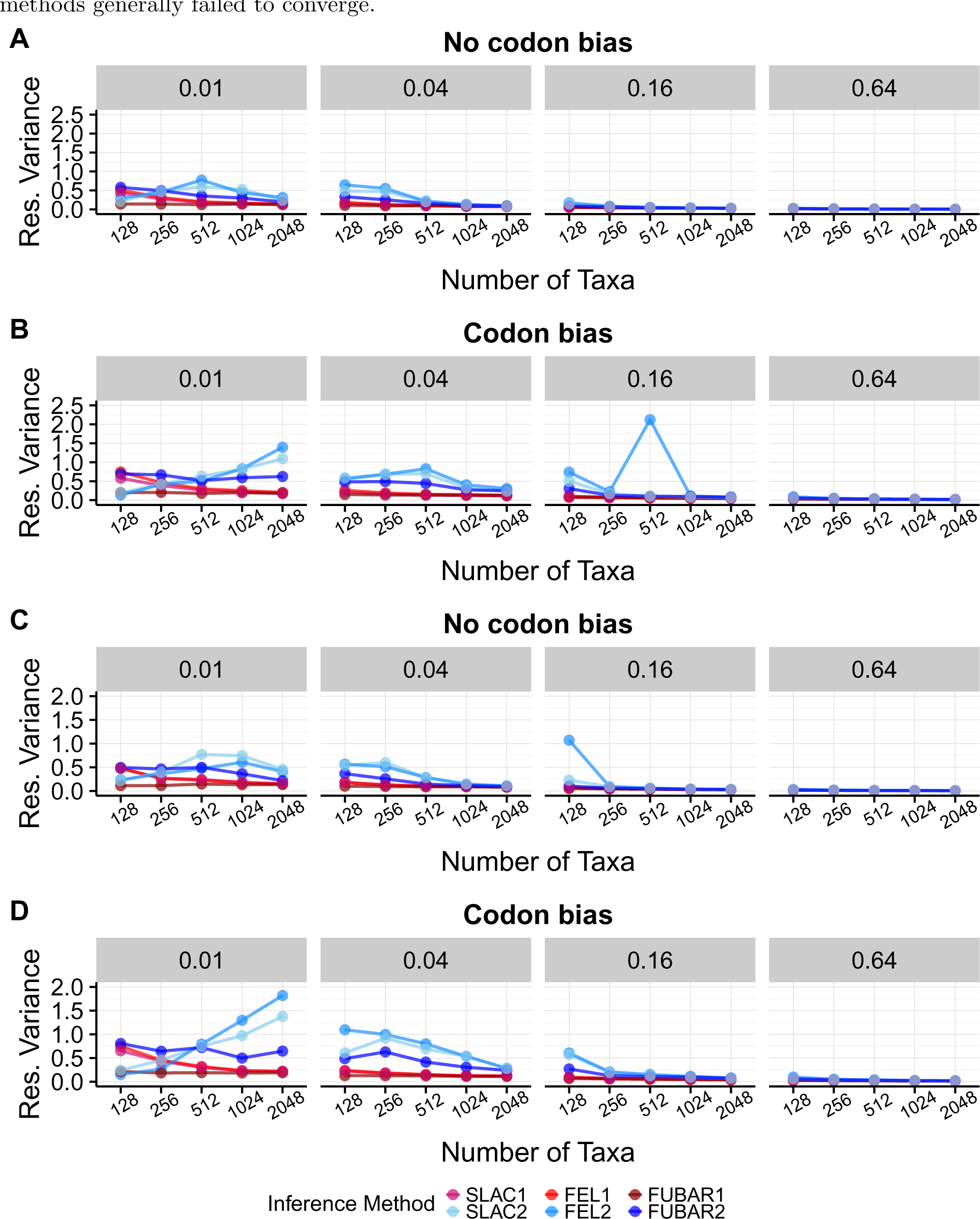
Variance in residuals for a linear model of inferred *dN/dS* regressed on true *dN/dS* across inference methods and simulation conditions. A) ∏_equal_ simulations with no codon bias. B) ∏_equal_ simulations with codon bias and equal nucleotide frequencies. C) ∏_unequal_ simulations with no codon bias. D) ∏_unequal_ simulations with codon bias. Note that certain FEL2 points are not present in the figure for conditions where the methods generally failed to converge.

**Figure S5.**
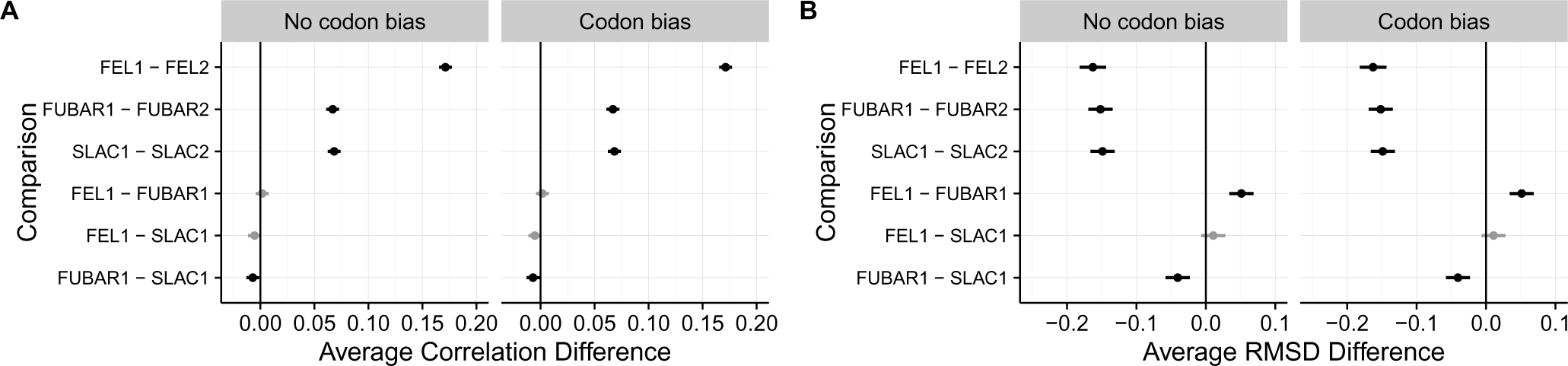
Pairwise comparisons of correlation strength and RMSD inference across methods, as determined through multiple comparisons tests. Results shown correspond to ∏_equal_ simulations. A) Results for multiple comparison tests of correlation strength. B) Results for multiple comparison tests of RMSD.

**Figure S6.**
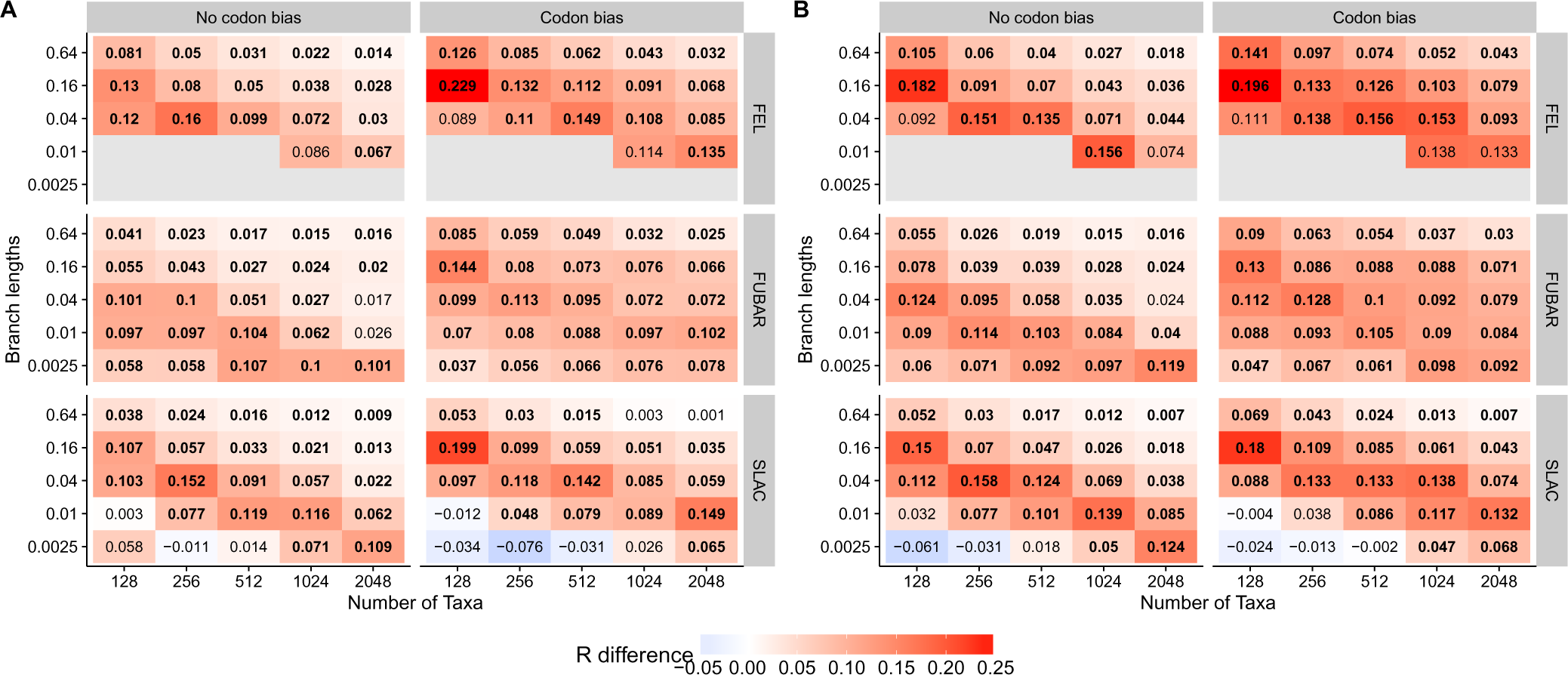
Correlation differences between one-rate and two-rate inference frameworks across simulation conditions, specifically for ∏_unequal_ simulations. Each cell indicates the average correlation increase from a two-rate to a one-rate platform, meaning that red cells represent conditions where one-rate methods outperform two-rate methods. Correlations were compared using paired *t-tests*, and significance was determined as Bonferroni-corrected *P <* 0.01. Bold values indicate that the difference is statistically significant. Conditions not shown (grey cells) represent cases for which fewer than five replicates converged, and thus comparisons were not performed.

**Figure S7.**
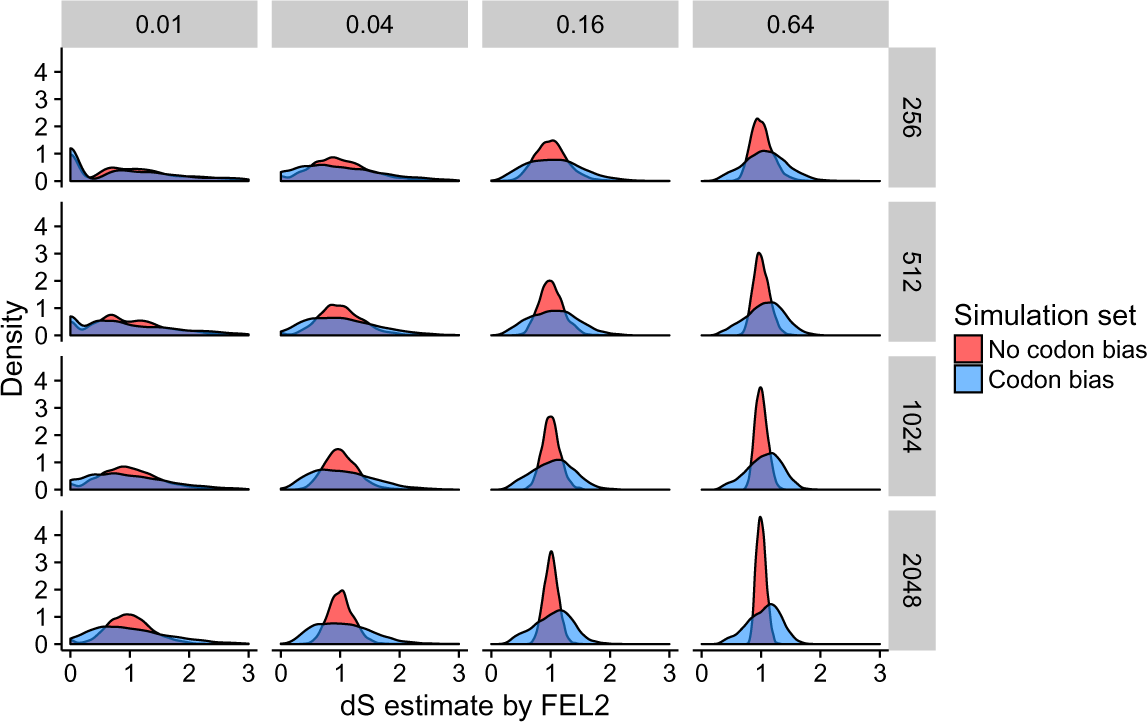
Inferred distributions of *dS* parameters by FEL2 for ∏_unequal_ simulations. Simulations with codon bias contained much more *dS* variation than did simulations without codon bias.

**Figure S8.**
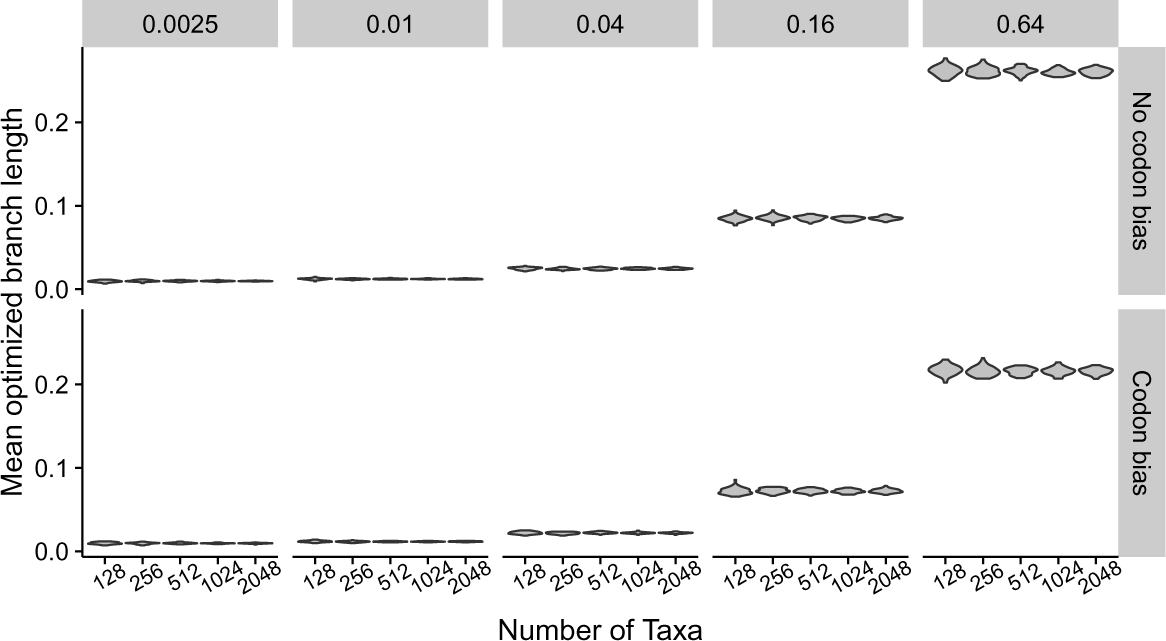
Optimized branch lengths across ∏_unequal_ simulations. For a given set of simulations using the same branch length, inferred distributions of optimized branch lengths were virtually identical.

**Figure S9.**
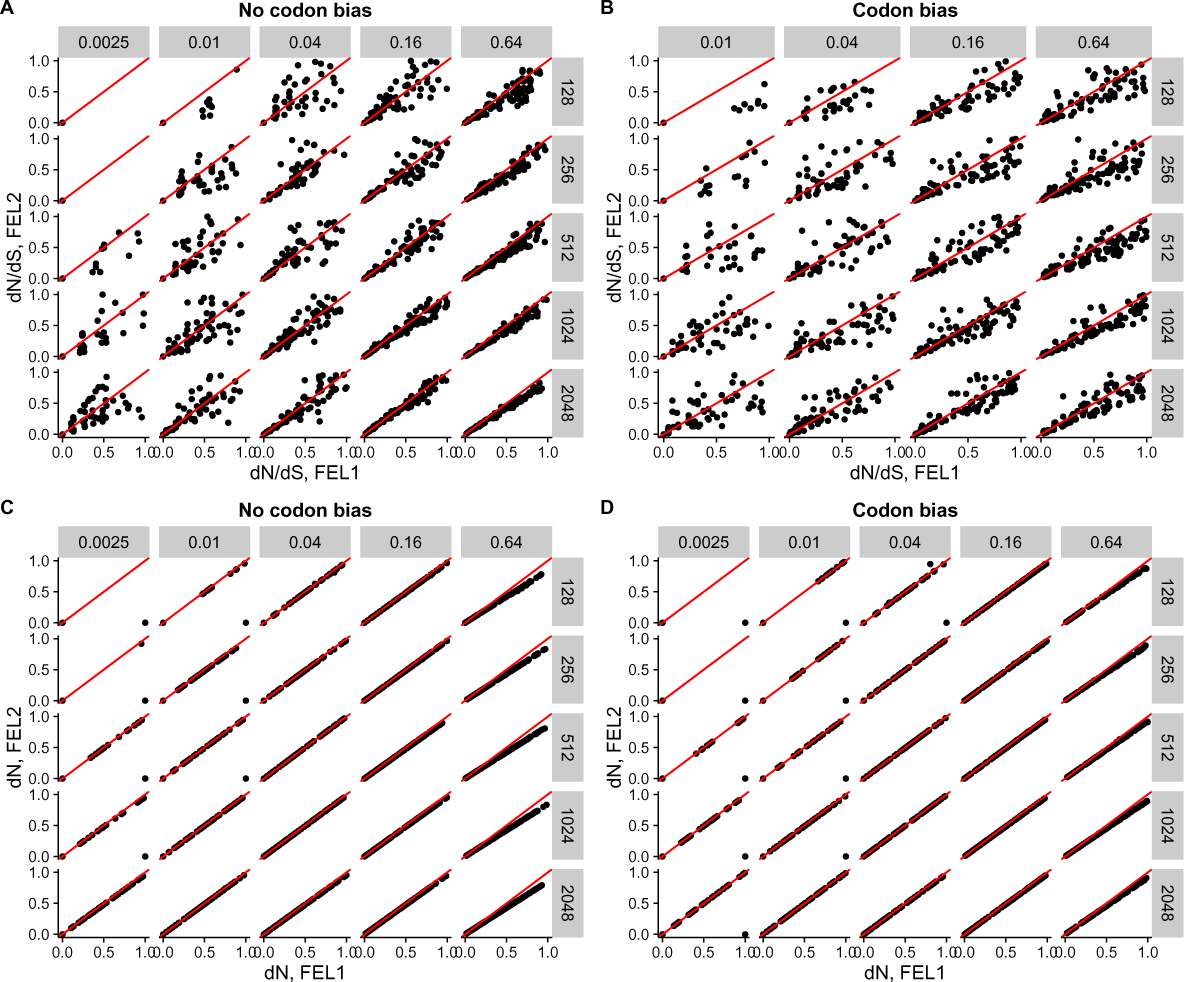
Direct comparison of *dN/dS* and *dN* estimates each between one-rate and two-rate frameworks. Results correspond to a single ∏_unequal_ simulation replicate, for each simulation condition. Each panel compares either the point *dN/dS* (A, B) or *dN* (C, D) estimate for each simulation condition, with branch lengths from left to right and number of taxa from top to bottom. X-axes indicate estimates with FEL1, and Y-axes indicate estimates with FEL2. The line in each panel is *x* =*y*.

**Figure S10.**
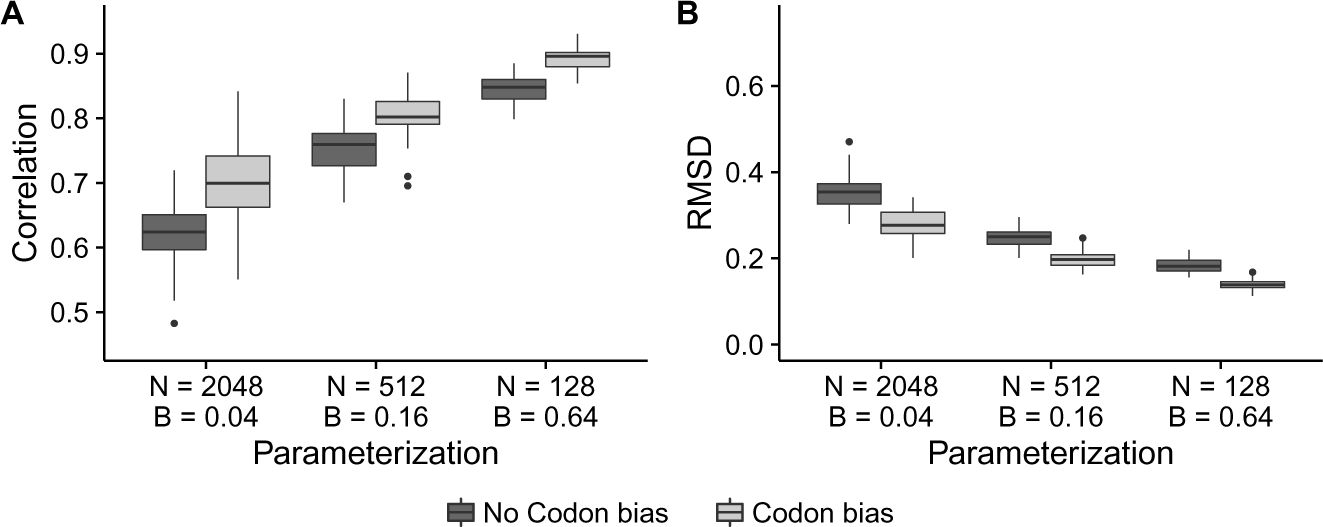
Divergence is more important than the number of sequences is for obtaining the equilibrium *dN/dS* value. Boxplots represent either A) correlation or B) RMSD, from FEL1 inference, across the 50 respective ∏_equal_ simulation replicates.

**Figure S11.**
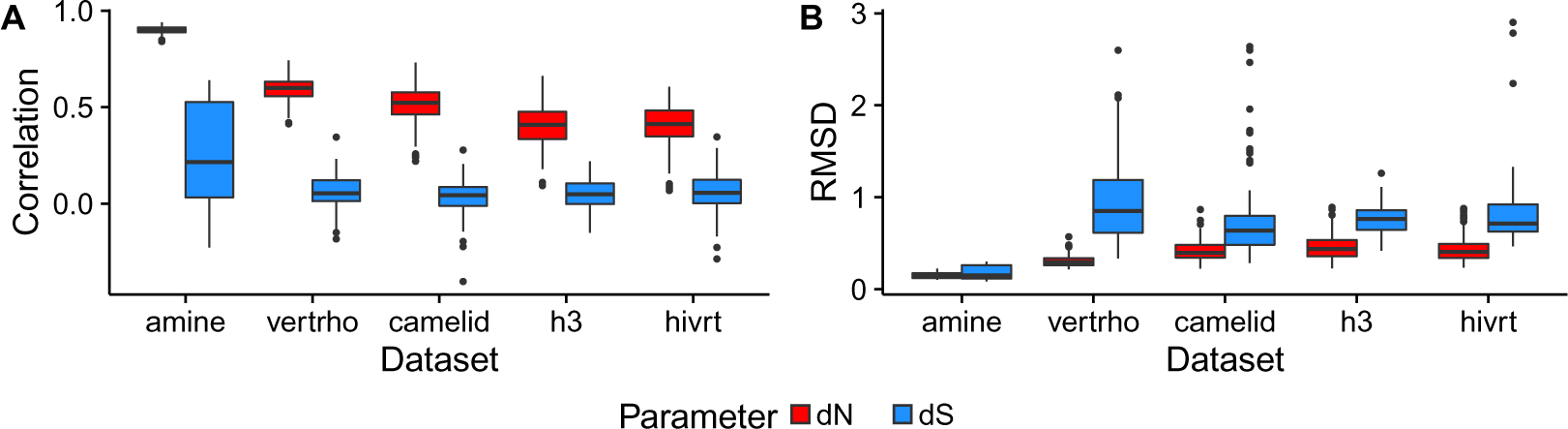
A) Correlation and B) RMSD for individual *dN* and *dS* parameter estimates with FUBAR2, for simulations with codon bias performed along empirical phylogenies.

